# Spatially-Resolved Live Cell Tagging and Isolation Using Protected Photoactivatable Cell Dyes

**DOI:** 10.1101/2020.02.28.966598

**Authors:** Alex S Genshaft, Carly G. K. Ziegler, Constantine N. Tzouanas, Benjamin E. Mead, Alex M. Jaeger, Andrew W. Navia, Ryan P. King, Tyler Jacks, Jeffrey F. Van Humbeck, Alex K. Shalek

## Abstract

Whether cultured *in vitro* or part of a complex tissue *in vivo*, a cell’s phenotype and function are significantly influenced by dynamic interactions with its microenvironment. To explicitly examine how a cell’s spatiotemporal activity impacts its behavior, we developed and validated a strategy termed SPACECAT—Spatially PhotoActivatable Color Encoded Cell Address Tags—to annotate, track, and isolate specific cells in a non-destructive, viability-preserving manner. In SPACECAT, a biological sample is immersed in a photocaged fluorescent molecule, and cells within a location of interest are labeled for further study by uncaging that molecule with user-patterned near-UV light. SPACECAT offers high spatial precision and temporal stability across diverse cell and tissue types, and is compatible with common downstream assays, including flow cytometry and single-cell RNA-Seq. Illustratively, we leveraged this approach in patient-derived intestinal organoids, a spatially complex system less amenable to genetic manipulations, to select for crypt-like regions enriched in stem-like and actively mitotic cells. Moreover, we demonstrate its applicability and utility on *ex vivo* tissue sections from four healthy organs and an autochthonous lung tumor model, uncovering spatially-biased gene expression patterns among immune cell subsets and identifying rare myeloid phenotypes enriched around tumor/healthy border regions. In sum, our method provides a minimally invasive and broadly applicable approach to link cellular spatiotemporal features and/or behavioral phenotypes with diverse downstream assays, enabling fundamental insights into the connections between tissue microenvironments and biological (dys)function.

## INTRODUCTION

From descriptions of gross anatomical features to characterization of subcellular organelles, modern *in situ* profiling methods have enabled insights into the organizational principles of biological structures. Next-generation single-cell genomic technologies hold great promise for new discoveries in health and disease, however they typically lack information regarding cellular location, morphology, microanatomy, and physiological context.

Current strategies for matching -omic measurements with spatial resolution rely on genetic models^1–5^, fixed or frozen samples^6–8^, or computational inference^9,10^. Approaches that rely on genetic modifications facilitate dynamic characterizations in model systems, but are not applicable to human clinical samples, and can be labor intensive to create and maintain. Methods reliant on fixed or frozen samples, meanwhile, offer static snapshots of complex systems and can be applied to nearly any tissue type, yet they are not compatible with all downstream measurements (e.g., biophysical assays) and offer limitations in some high-throughput genomic assays (e.g., due to RNA degradation or input cell number). Similarly, computational methods rely on *a priori* knowledge of sample structure, which is possible in many healthy tissues but cannot be assumed in novel biological systems or heterogeneous clinical samples.

To address these shortcomings, we introduce SPACECAT (Spatially PhotoActivatable Color Encoded Cellular Address Tags), a method for tagging and tracking cells from user-selected regions, enriching for physically observable features, and isolating cells for arbitrary downstream processing. We designed SPACECAT to be minimally invasive and compatible with a diverse set of input samples, and to yield live cells that have been tagged in a user-designated fashion encoding prior tissue location, morphology, neighborhood, and/or other observable behaviors. In SPACECAT, a biological sample is immersed in a photocaged dye and specific areas of interest are illuminated with a spatially controlled near-UV light source. The near-UV light uncages the dye, generating a fluorescent molecule *in situ* that stains cells within the photoactivated regions. These fluorescent cells can then be tracked over time and isolated via flow sorting, enabling arbitrary combination of imaging modalities with biophysical, functional, or genomic assays, all while retaining registry of user-tagged cells.

Here, we apply SPACECAT to 2D *in vitro* cultures, 3D human intestinal organoids, and mouse tissue sections across health and disease. We validate its spatial selectivity and precision through comparisons with known structural phenotypes, and apply it to a lung tumor mouse model to uncover previously unknown spatial heterogeneity. Crucially, SPACECAT can be incorporated into pre-existing sample processing pipelines, does not require complex chemistry to be performed upon samples, and can be applied to samples across donor, species, or tissue of origin. Thus, SPACECAT holds promise to link microanatomical spatial knowledge and dynamic cellular phenotypes with the unbiased molecular resolution of highly parallel single-cell assays.

## RESULTS

### Nitroveratryloxycarbonyl caging of carboxylate groups enables rapid, stable, spatially selective generation of fluorescent calcein within live cells

To arbitrarily tag live cells for tracking and isolation, we chose calcein as a scaffold to modify with nitroveratryloxycarbonyl (NVOC) photolyzable protecting groups. As a commonly used viability dye^11^, calcein enables determination of cell health as part of our cell tagging scheme without occupying an additional fluorescence channel. Commercially available calcein AM has seven protected carboxylic acids, all of which must be deprotected for maximal fluorescence^12^. Six of these acids are protected as an acetoxymethyl esters, and the last is protected as a lactone. Upon entering the cell, the acids are deprotected via esterase cleavage and the molecule becomes fluorescent; the net negative charge that amasses during deprotection is thought to retain the fluorescent molecule in the cell (**Supplementary Fig. 5b)**.

Here, we photocaged calcein’s EDTA-like moieties by adapting a previously described synthetic schema for creating a xanthene scaffold that is specifically functionalized at the 4’ and 5’ positions (**Fig. 1a**, see **Methods**)^13^. First, the xanthene scaffold (**compound 1, Supplementary Fig. 1**) was synthesized with methyl groups in the 4’ and 5’ positions and acetoxymethyl ester protecting groups on the two alcohols with a 50% yield over two steps. This molecule was then brominated to form **compound 2** with a 68% yield (**Supplementary Fig. 2**). In parallel, the NVOC protected EDTA-like moiety (**compound 3, Supplementary Fig. 3**) was synthesized by combining 2-nitro-4,5-dimethoxybenzyl bromide and *N*-Boc-iminodiacetic acid, removing excess bromide with a silica plug, and finally removing the Boc protecting group in trifluoroacetic acid with 91% yield. Lastly, **compounds 2** & **3** were combined in the presence of sodium iodide and 1,8 bis(dimethylamino)naphthalene (proton sponge) in dry acetonitrile until the reaction went to completion. The final **compound 4**, which we term calcein NVOC, was purified using a preparative TLC plate with 28% yield including a small amount of residual solvent (**Fig. 1a, Supplementary Fig. 4**).

**Figure 1.**
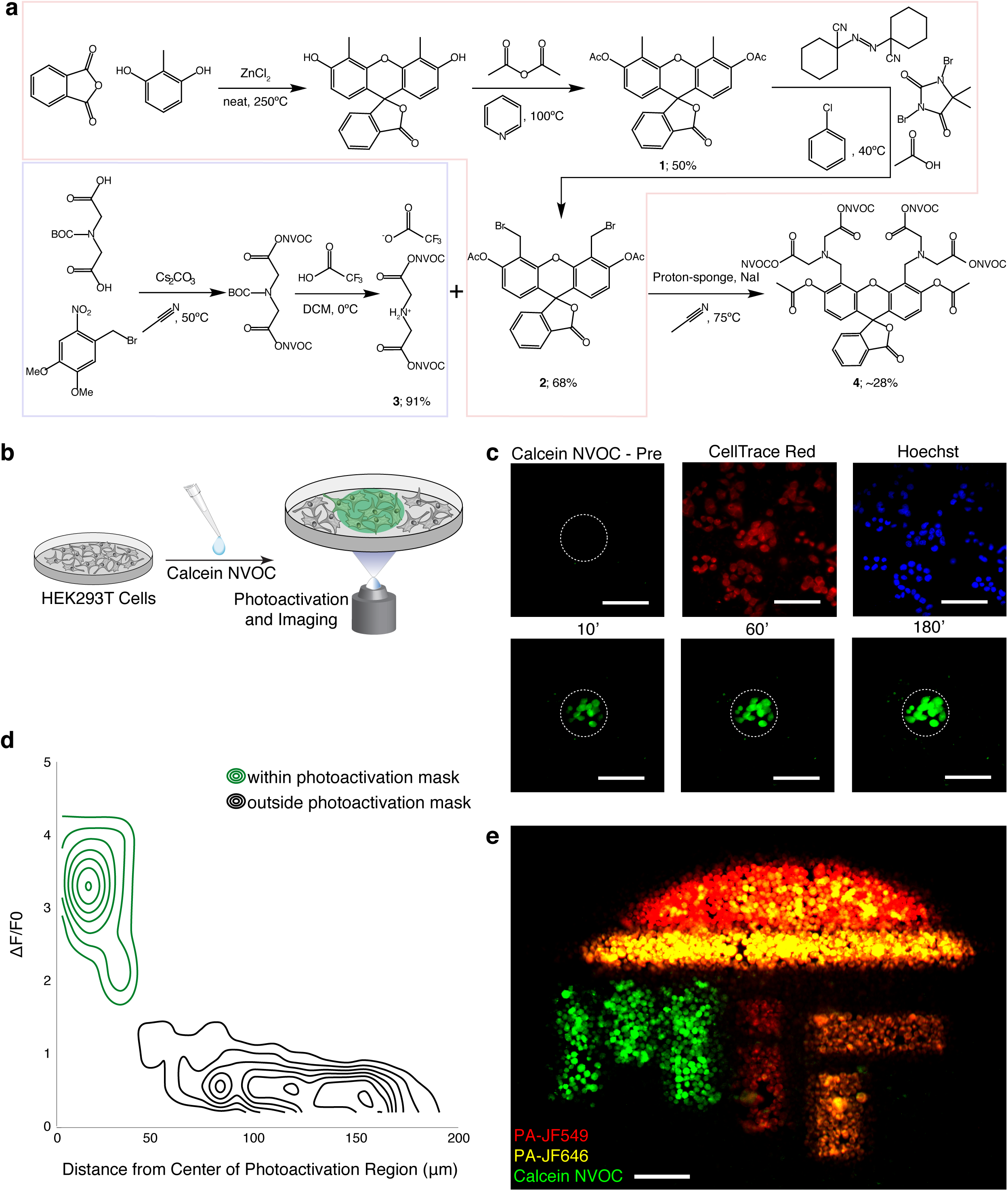
The SPACECAT protocol enables spatially precise, temporally stable fluorescence signals in arbitrary regions of interest. **a**. Synthetic schema for calcein NVOC (see **Methods**). **b**. HEK293T cells were stained with calcein NVOC, and imaged before photoactivation and afterwards for three hours. **c**. Representative time course images from a single field of view. Presented images of calcein NVOC were taken before photoactivation (“calcein NVOC Pre”), and 10 minutes after (10’), 60 minutes after (60’), and 180 minutes (180’) after photoactivation. Images of CellTrace Red and Hoechst stains were taken 180 minutes after photoactivation. Dashed line: photoactivated region; scale bar = 100μm. **d**. Contour plots representing the spatial distribution of fluorescence changes in photoactivated regions (green lines, n = 68 cells across 8 fields of view) and regions outside of photoactivation mask (black lines, n = 1,347 cells across 8 fields of view). ΔF/F_0_ is defined as a cell’s change in mean fluorescence 180 minutes after photoactivation, divided by the cell’s pre-photoactivation mean fluorescence. **e**. Representative composite image of HEK293T cells stained and photo-uncaged within the “MIT” logo region. By sequential addition of 3 photoactivatable probes (calcein NVOC, PA-JF549, PA-JF646) and leveraging different photoactivation thresholds (10 seconds for calcein NVOC, 0.5 seconds for PA-JF549 and PA-JF646), we achieve 5-color encoding. Scale bar = 100μm. Green: uncaged calcein NVOC; yellow: uncaged PA-JF549, red: uncaged PA-JF646.

To test calcein NVOC, we applied it to HEK293T cells in complete media and photoactivated user-defined regions of interest using a digital micromirror device (**Fig. 1b**). Images taken before and after photoactivation revealed significant fluorescence increases, measurable immediately after active illumination and stable throughout the 3-hour duration of this time course experiment (**Fig. 1c, Supplementary Fig. 5c**). This temporal stability is critical for downstream experimental flexibility, enabling sample dissociation protocols, flow cytometry, and even subsequent functional or biophysical measurements following cell tagging. Alternately, cells can be tracked *in situ*, enabling experiments that combine behavioral readouts with spatial location and downstream measurements.

Moreover, fluorescence increases were specific to photoactivated regions of interest. This was quantified by comparing the change in fluorescence inside versus outside the photoactivated region (**Fig. 1c, 1d**; p-value < 0.0001, Student’s two-sample t test of 68 cells within vs. 1347 outside the photoactivated region; **Methods**; **Supplementary Fig. 5c**). Finally, to demonstrate the spatial control of SPACECAT’s tagging procedure, we photoactivated HEK293T cells within a mask of MIT’s logo following treatment with calcein NVOC as well as two photoactivatable dyes^14^ with distinct fluorescence spectra attached to an NHS-moiety (PA-JF549, PA-JF646, **Fig. 1e, Supplementary Fig. 6a**). Taking advantage of the different thresholds for photoactivation between distinct dyes, we uniquely tagged cells with 5 distinct color combinations, and confirmed fluorescence was maintained through dissociation and flow cytometry (**Supplementary Fig. 6b-i**). This demonstrates the multiplexing capacity of calcein NVOC and an approach that enables simultaneous encoding of distinct microanatomical neighborhoods.

### Flow cytometry distinguishes calcein NVOC tagged cells with high sensitivity

To demonstrate the specificity of both tagging and isolation of cells from desired regions, we performed a species-mixing experiment where HEK293T (human) and NIH/3T3 (mouse) cells were co-cultured in a spatially structured manner (**Supplementary Fig. 7a**). After incubation with calcein NVOC, one of the two zones of cells (i.e., either mouse or human) was photoactivated, creating three conditions: 1) co-cultured NIH/3T3 and HEK293T, with tagged NIH/3T3, 2) co-cultured NIH/3T3 and HEK293T, with tagged HEK293T, and 3) co-cultured NIH/3T3 and HEK293T, both un-tagged. Cells were isolated by fluorescence-activated cell sorting (FACS), and calcein NVOC+ fluorescent cells from each experimental condition were individually sorted for single-cell RNA-Seq (scRNA-seq; **Supplementary Fig. 7b; Methods**)^15,16^. Single cell libraries were aligned to a joint human and mouse reference genome, which identified each cell’s species of origin (**Supplementary Fig. 7c**). Of the 48 cells passing quality thresholds from the HEK293T tagged condition, 45 aligned to the human transcriptome; from the NIH/3T3 tagged condition, 61 cells passed quality thresholds, 60 of which aligned to the mouse transcriptome. When compared with the unphotoactivated control, these distributions were significantly enriched for the desired species (p-values = 0.026 and 5.5e-19, for human and mouse, respectively, Fisher’s exact test). Collectively, these data demonstrate that the photoactivated calcein NVOC fluorescence signal persists through dissociation and can be detected by flow cytometry, demonstrating SPACECAT’s utility for tagging and extracting cells.

### Precise cell tagging of human intestinal organoid morphological features confirms stem-like program in budding crypts

To test SPACECAT’s capacity to tag specific spatial features within a complex 3D biological structure in a system where genetic transformation is difficult, we applied SPACECAT to enrich for known morphological features within human intestinal organoids^17^. Human organoids are highly structured and possess significant cellular heterogeneity, with crypt-like protrusions known to be enriched for highly-proliferative, stem-like populations. To isolate cells from crypt regions, we photoactivated protrusions on the organoid periphery, producing fluorescence signals that were spatially specific and did not include cells in core or villi regions of organoids (**Fig. 2a, 2b**). Organoids were dissociated, and populations of calcein NVOC+ cells and background (i.e., unphotoactivated) cells were isolated by FACS. Subsequent scRNA-seq using the Seq-Well platform^18–20^ revealed three cell states within this organoid model (**Fig. 2c, Supplementary Table 1**), and SPACECAT enabled us to map cells to their microanatomical origin (**Fig. 2d**). Using previously described gene lists^21^, we scored each cell and categorized them into various stages of the cell cycle (G1, S, and G2/M; **Fig. 2e**). We found that a significantly larger proportion of cells derived from the protruding crypt regions had entered the cell cycle compared to non-crypt regions (**Fig. 2f**; p-value < 0.001, Chi-squared test). Calcein NVOC+ cells were also significantly enriched for stem-like populations^17^, while the bulk organoid consisted of roughly equal proportions of stem-like cells and mature enterocytes with a small fraction of developing goblet cells (**Fig. 2g, 2h**; p-value < 0.001, Chi-squared test). Our findings align with previous knowledge of intestinal organoid structure, as crypts are widely accepted as the main site of intestinal proliferation and harbor the intestinal stem cell niche^17^. Thus, SPACECAT affords a simple strategy for selecting, labeling, and enriching cells from user-specified regions within 3D human organoids.

**Figure 2.**
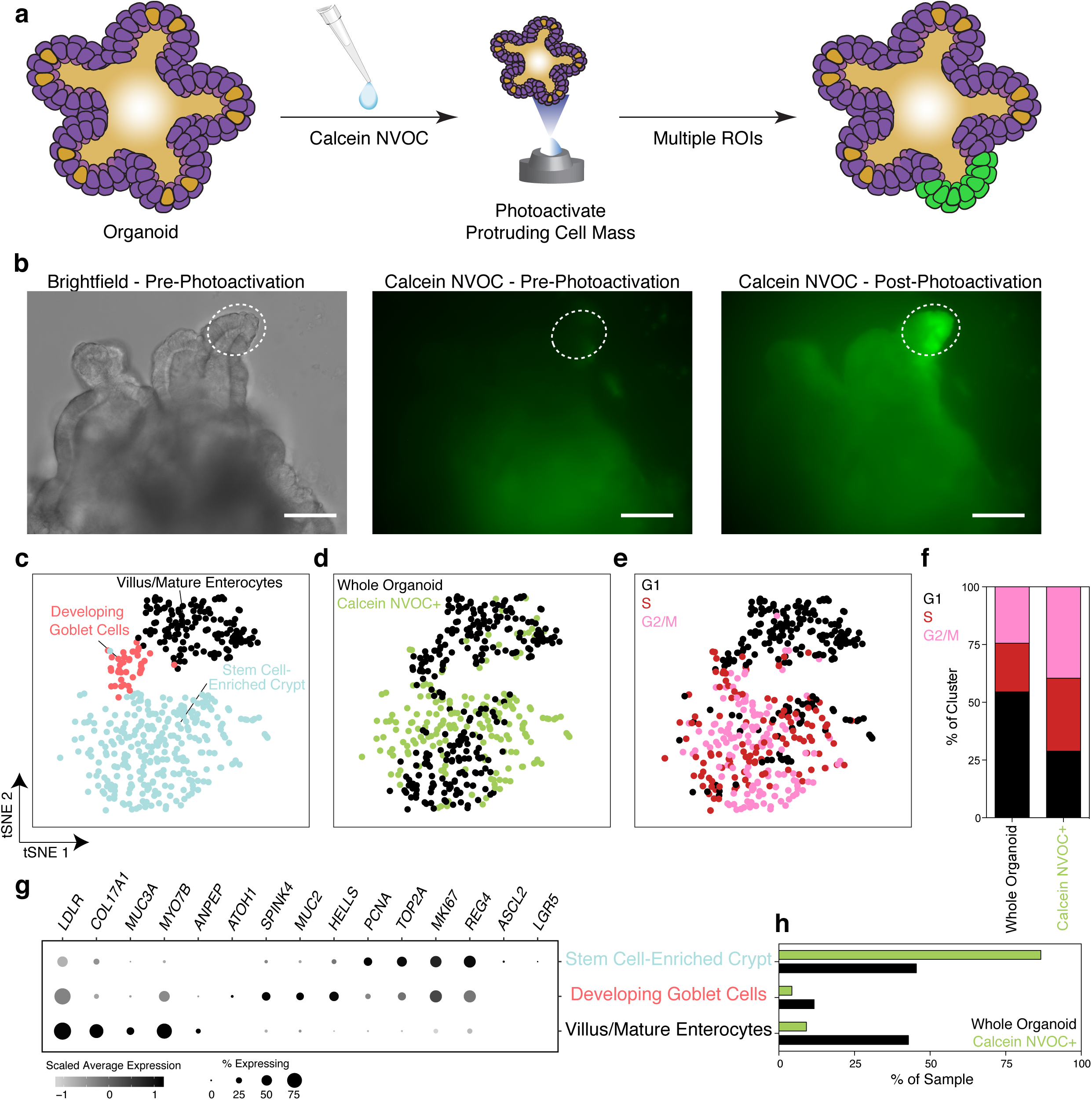
Application of SPACECAT for physical isolation of stem cell niches from human small intestinal organoid models. **a.** Schematic of organoid crypt selection with the SPACECAT protocol. **b.** Representative images of cultured organoids and example selected regions for photoactivation and selection. Dashed line: photoactivated region; scale bar = 100μm. **c.**-**e.** tSNE of 453 single cells from whole organoid and sorted photoactivated green regions. **c.** cells colored by Louvain clusters; **d.** cells colored by cell cycle (black: G1, red: S, pink: G2/M); **e.** cells colored by experimental condition (whole organoid: black; photoactivated region: green). **f.** Cell cycle stage by experimental condition. p-value<0.001 by Chi-square test (Calcein NVOC+: 187 cells, Whole Organoid: 266 cells). **g.** Selected differentially expressed genes marking each Louvain cluster. Circle size denotes percent of each cluster expressing a given gene, circle color denotes relative digital gene expression compared to other clusters. **h**. Cluster membership by experimental condition. p-value<0.001 by Chi-square test.

### Calcein NVOC is compatible with diverse tissues

To demonstrate the versatility of SPACECAT in complex tissue microenvironments, we photoactivated arbitrary regions in live tissue slices of mouse brain, small intestine, lung, and spleen (**Fig. 3a-d, Supplementary Fig. 8a**). Despite the range of cell types and tissue structures (e.g. extracellular matrix types and composition) within these tissues, photoactivation of calcein NVOC provided strong fluorescence signals that were specific to the desired regions. Critically, the SPACECAT protocol required no alterations or optimization between distinct tissue types, demonstrating wide applicability and compatibility.

**Figure 3.**
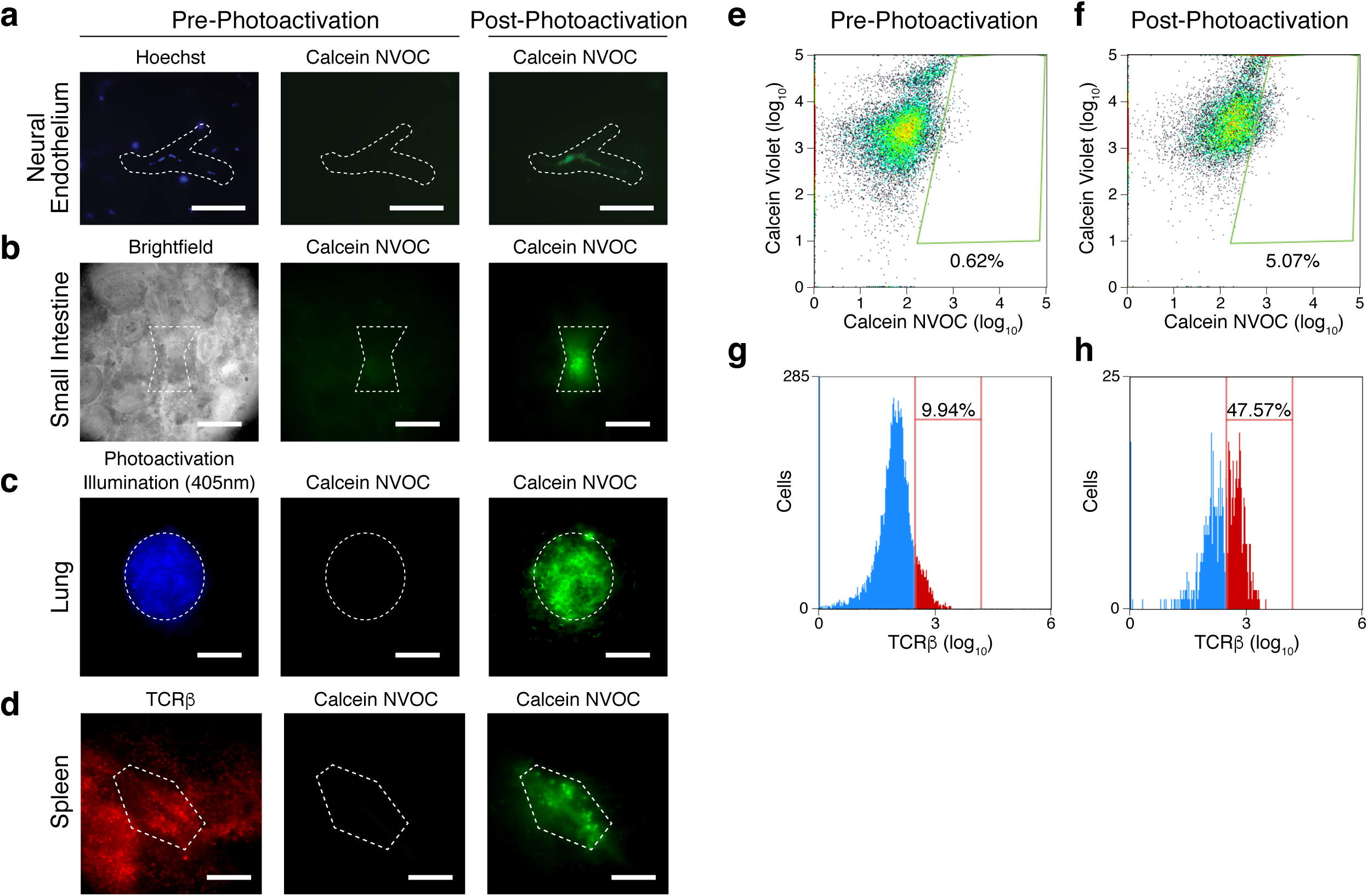
SPACECAT enables tagging of precise regions in live tissue sections. **a.**-**c.** Various mouse tissue samples were sectioned live, stained with calcein NVOC, and photoactivated in arbitrary regions (white dashed regions). **a**. endothelial cells within brain tissue (scale bar = 100μm), **b.** jejunum (scale bar = 50μm), **c.** lung (scale bar = 50μm). **d.** Representative images of mouse spleen, T cell rich areas were selected for photo-tagging (dashed white region). Left: TCRβ chain antibody (red); middle: calcein NVOC prior to photoactivation; right: calcein NVOC after photoactivation. Scale bar = 50μm. **e.** and **f.** Flow cytometry of calcein NVOC (x-axis, marker of viability and photoactivation) vs. calcein violet (y-axis, marker of viability) in un-photoactivated (**e.**) and photoactivated (**f**.) splenic slices of T-cell rich areas. **g.** and **h.** Abundance of TCRβ positive cells among all splenic cells (**g.**) and photoactivated cells (**h.**) from photoactivated splenic slices.

Next, to test the precision of SPACECAT cell tagging within a heterogeneous tissue, as well as the ability for calcein NVOC fluorescence signal to be retained after tissue dissociation, we photoactivated T cell-rich regions found in close proximity to vasculature within the spleen (likely representing periarteriolar lymphoid sheaths; **Fig. 3d, Supplementary Fig. 8a**)^22^. As expected, photoactivation of calcein NVOC produced fluorescent cell populations that were not present before photoactivation, as assessed by both imaging and flow cytometry (**Fig. 3e, 3f**). Further, when analyzed by flow cytometry, the fluorescent cell population was ∼5x enriched for cells positive for anti-TCR β chain antibody, supporting successful isolation of regions of interest (**Fig. 3g, 3h**). Notably, while not directly tested in this experiment, we began photoactivation of spleen sections 7 hours before signal was detected by flow cytometry, indicating a potentially much longer period of fluorescence signal stability than shown in our *in vitro* time course (**Fig. 1c, Supplementary Fig. 5c**). Thus, the SPACECAT protocol enables enrichment and multimodal profiling of cells from user-specified spatial regions in diverse tissue architectures and is easily integrated in complex sample processing pipelines.

### User-directed cell tagging uncovers spatial heterogeneity in tumor infiltrating myeloid cells

Finally, we applied SPACECAT to a genetically engineered mouse model of lung adenocarcinoma that phenotypically recapitulates key aspects of human disease. In the Kras-p53 (Kras^LSL-G12D/+^, p53^fl/fl^; KP) model^23^, intratracheal administration of adenovirus expressing Cre recombinase induces oncogenic *Kras* expression with simultaneous deletion of p53, leading to transformation of alveolar type II cells in the context of the healthy lung stroma and natural immune response. To test SPACECAT in the KP model, we dissected whole tumors 16 weeks post initiation from the lungs of two mice, and interrogated the tumor periphery and core by exposing intact tumor sections to calcein NVOC, photoactivating desired regions, sorting cells by calcein NVOC fluorescence, and generating single cell RNA-seq libraries using Seq-Well (**Fig. 4a, Supplementary Fig. 8b, 8c**).

**Figure 4.**
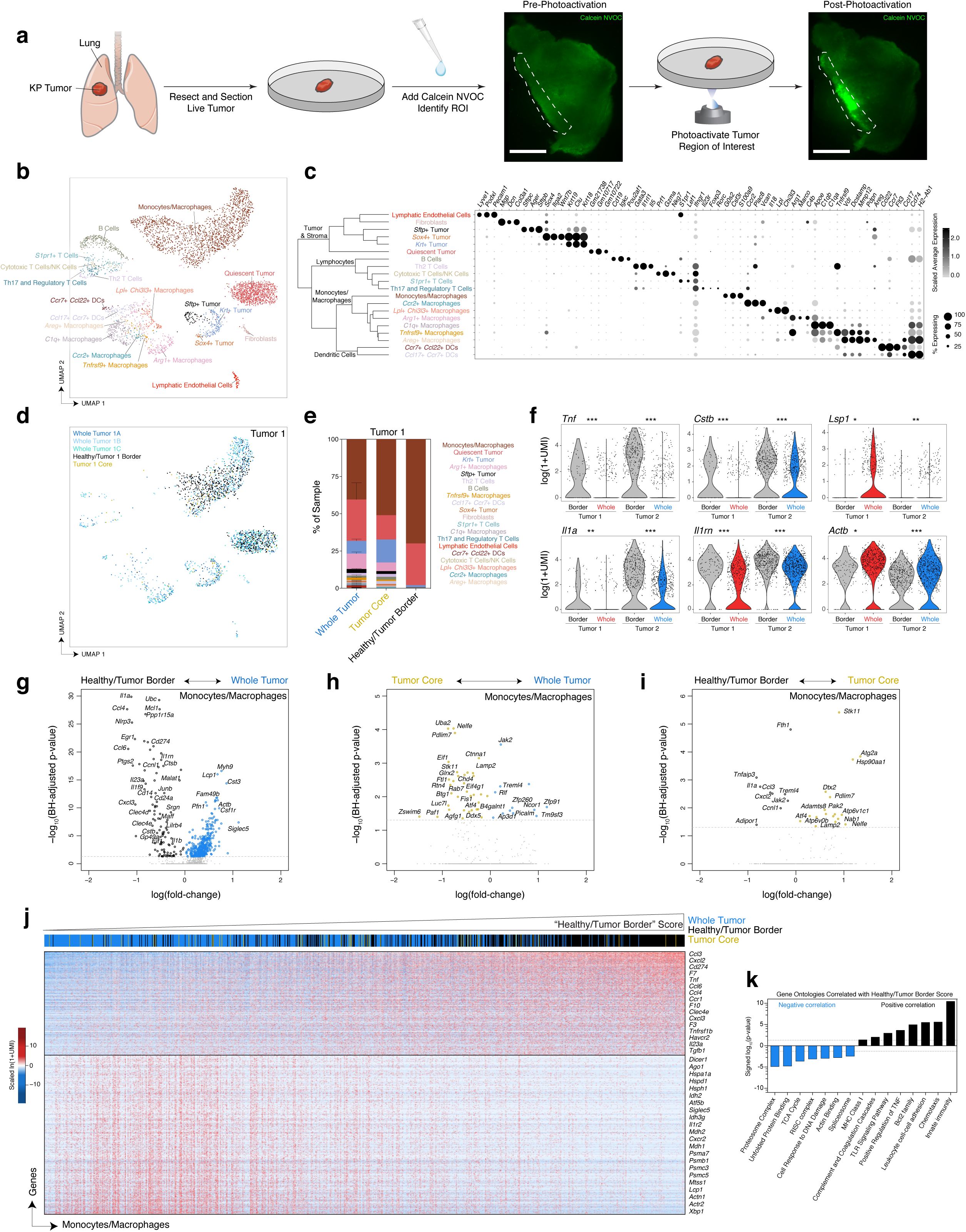
Applying SPACECAT to a KP autochthonous lung tumors uncovers spatial heterogeneity of immune infiltrate. **a.** Experimental schematic and representative stitched images of KP lung tumors slices in FITC fluorescence channel before photoactivation (left) and after photoactivation of regions within the white dashed lines (right). Scale bar = 500μm. **b.** UMAP embedding and Louvain clusters of 4,447 single cells, annotated by cell type and state. **c.** Hierarchical clustering of cell types and dot plot of top differentially expressed genes between each cluster and all other cells. All genes significant by Bonferroni adjusted p-value < 0.001. Dot color represents average expression per cluster, dot size represents percent expression per cluster. **d.** UMAP embedding with points corresponding to cells only in Tumor 1. Points colored by sort condition, shades of blue represent triplicate Seq-Well arrays from the Whole Tumor. Black points represent single cells photoactivated and sorted from the Healthy/Tumor Border. Yellow points represent single cells photoactivated and sorted from the Tumor Core. **e**. Relative cell type compositions by tumor region in Tumor 1. Error bars indicate mean ± SEM. **f.** Expression of individual genes among Monocyte/Macrophage cells by tumor region in Tumor 1 (Healthy/Tumor Border vs. Whole Tumor). Bonferroni-adjusted p-value < 0.001 ***, < 0.01 **, < 0.05 *. **g.-i.** Volcano plots of differentially expressed genes between Monocyte/Macrophage cells by tumor region in Tumor 1, dashed lines correspond to p-value = 0.05. **g.** Healthy/Tumor Border vs. Whole Tumor, **h.** Tumor Core vs. Whole Tumor, **i.** Healthy/Tumor Border vs. Tumor Core. Horizontal dashed line represents Bonferroni-adjusted p-value<0.05. Whole Tumor: 674 cells, Healthy/Tumor Border: 480 cells, Tumor Core: 53 cells. **j.** Heatmap of genes significantly correlated with Heatlhy/Tumor Border Score among Tumor 1 Monocyte/Macrophage cells (blue: lower relative expression, red: higher relative expression). Cells are ordered by expression of Healthy/Tumor Border Score (color bar represents original location of each cell: blue, Whole Tumor; black, Healthy/Tumor Border; yellow, Tumor Core). All genes significant by p-value<0.01. **k.** p-values of significantly enriched gene ontologies among genes positively correlated with Healthy/Tumor Border Score (black bars) and genes negatively correlated with Healthy/Tumor Border Score (blue bars). Dashed lines correspond to p-value = 0.05.

After preprocessing and filtering for high-quality cells, we recovered 4,447 single cells and 27,923 unique genes from two tumors (**Supplementary Table 2**). For cell type identification, we reduced this high-dimensional data into a lower-dimensional manifold using principal component analysis (PCA) over variable genes, clustered cells using a mutual k nearest-neighbor graph, and visualized these clusters on a UMAP embedding (**Fig. 4b; Methods**)^24^. This approach uncovered 20 distinct cell types representing multiple known lymphoid, myeloid, and stromal populations associated with the KP tumor model, including 4 distinct subpopulations of tumor cells, termed *Sox4+* Tumor, *Krt+* Tumor (defined by upregulation of multiple keratin genes including *Krt19, Krt18, Krt7*), *Sftp+* Tumor (defined by upregulation of surfactant genes including *Sftpb, Sftpc, Sftpd*), and a Quiescent Tumor cell type (**Fig. 4c, Supplementary Table 3**). Notably, we recovered 9 populations of monocytes, macrophages, and dendritic cells represented in both tumors. We termed the largest subpopulation of myeloid cells “Monocytes/Macrophages,” defined by expression of well-described markers of monocytes and activated macrophages including *Csf3r, G0s2, Cd14*, and *Fcgr3* as well as multiple inflammatory cytokines including *Tnf, Ccl4, Cxcl3, Il1b, Il1a* and *Ccl3*. The “Monocytes/Macrophages” cell cluster is distinct from the 6 other populations of macrophages, which each express a more restricted and mutually-exclusive set of functional genes, suggesting greater differentiation and specialization (**Supplementary Table 3**). To our knowledge, this level of cellular and spatial resolution of myeloid populations within the KP model has not been described previously.

All cell types were recovered from both Tumor 1 and Tumor 2, however at different relative abundances (**Fig. 4d, 4e**: Tumor 1; **Supplementary Fig. 9a, 9b**: Tumor 2). Notably, in Tumor 2, we recovered a much higher proportion of B cells, T cells, and dendritic cells, potentially indicating either a more immune-permissive environment within Tumor 2, or that we may have captured a tumor-associated tertiary lymphoid structure. To control for tumor-intrinsic heterogeneity, we compared the abundance of cell types between calcein NVOC+ Healthy/Tumor Border sorted cells and cells from the unsorted, Whole Tumor arrays in each tumor separately (**Fig. 4e, Supplementary Fig. 9b**). In both Tumor 1 and Tumor 2, there were significant differences in the abundance of the Monocyte/Macrophage population between the Healthy/Tumor Border-derived cells and the unsorted Whole Tumor (p-value <0.05 by Student’s t-test, both comparisons). In addition, we observed a significant expansion of *C1q*+ Macrophages in the Tumor/Healthy Border of Tumor 1 (p-value <0.05 by Student’s t-test) – this cell type was defined by elevated expression of complement components *C1qa, C1qb*, and *C1qc*, cathepsins *Ctsh, Ctsc*, and *Ctss*, as well as *Cx3cr1*, which is involved with macrophage migration to inflamed sites. Notably, in Tumor 1, we observed that the Tumor Core samples recovered many of the cell types present within the Whole Tumor, including diverse lymphocyte populations, specialized myeloid cells, and each tumor subtype. Conversely, in the Tumor 1 Healthy/Tumor Border sample, the majority of the cells were restricted to the Quiescent Tumor or Macrophage/Monocyte cell types. Together, this data supports profound spatial heterogeneity among tumor subtypes and infiltrating lymphocytes within the KP tumor model.

Given significant changes in the relative abundance of Monocyte/Macrophage between the Healthy/Tumor border and Whole Tumor samples from both tumors, we tested for differentially expressed genes by location within this cell type. From Tumor 1, we observed 117 genes downregulated and 372 genes upregulated in the Whole Tumor compared to the Healthy/Tumor Border (**Fig. 4g, Supplementary Table 3**); from Tumor 2, we observed 31 downregulated and 80 upregulated genes in the same comparison (**Supplementary Fig. 9c, Supplementary Table 3**). Genes *Tnf, Il1a, Il1rn*, and *Ctsb* were consistently expressed at higher levels in the Healthy/Tumor Border-derived monocytes and macrophages, suggesting inflammatory macrophages may have limited access to the tumor core or they represent recently arrived cells that still retain pro-inflammatory functions (**Fig. 4f**). Moreover, we observe genes associated with transendothelial migration and motility, including *Lsp1* and *Actb*, consistently downregulated among Monocytes/Macrophages in calcein NVOC+ tagged Healthy/Tumor Border cells, potentially signifying a limited capacity for migration into deep avascular tumor spaces among myeloid cells found at the tumor border (NB: we confirmed metrics of cell quality and complexity, UMI/cell and unique genes/cell, were identical between all groups compared in **Fig. 4f**)^25^. In line with these findings, a direct comparison between Monocytes/Macrophages found in the Tumor Core vs. those found at the Healthy/Tumor Border identified *Tnf, Il1a, Ccl3, Cxcl2, Jak2*, and other inflammatory markers similarly upregulated among Monocytes/Macrophages restricted to the tumor periphery. Conversely, we found genes involved in autophagy and phagolysosome formation (*Lamp2, Atg2a, Rab7, Atp6v1c1, Atp6v0b*) upregulated among the Tumor Core resident Monocytes/Macrophages in both comparisons, as well as upregulation of disintegrin and metallopeptidase *Adamts8*, which may aid in cellular access to deep tumor regions (**Figure 4h, 4i, Supplementary Table 3**).

Using the 117 genes with increased expression among Monocytes/Macrophages at the Healthy/Tumor Border in Tumor 1, we scored all cells from Tumor 1 belonging to this cluster by their expression of this set of genes. We then ranked cells according to this “tumor periphery” score, akin to methods for pseudo-temporal ordering of cells, allowing us to pseudo-spatially order all Monocyte/Macrophage cells found within the Healthy/Tumor Border, Whole Tumor, and Tumor Core samples from Tumor 1 (**Fig. 4j**). We assessed which cellular pathways were associated with this ranking and found 887 genes negatively correlated and 577 genes positively correlated with a Healthy/Tumor Border phenotype (**Supplementary Table 3**). Among the gene expression programs that increase as monocytes and macrophages take on a more “healthy/tumor border-like” phenotype, we found significant enrichment for chemokines and cytokines involved in monocyte, neutrophil, and lymphocyte recruitment, including *Ccl1, Ccl3, Cxcl3, Cxcl2, Ccl4, Ccl6, Il1a*, and *Il1b*, as well as multiple toll-like receptor genes (*Tlr4, Tlr2, Tlr6, Tlr13)* and genes involved in inflammasome formation, such as *Nlrp3* (**Fig. 4j, 4k**). We also observed upregulation of coagulation factors *F3, F7, F10*, and *Plau*, and peptidases *Adam8, Tpp1, Furin*, and *Npepps*, suggesting tumor-periphery associated macrophages are mounting a wound-repair response and actively remodeling the local vasculature and lung stroma. Notably, among gene programs significantly negatively correlated with location at the tumor periphery, we found differential expression of *Cxcr2*, which has been previously associated with deep-tumor infiltrating tumor-associated macrophages and an anti-inflammatory phenotype^26^. In the “non-periphery-like” cells we also found upregulation of genes involved in protein homeostasis (*Hspa1a, Hspe1, Dnajc3, Hsp90ab1*), RNA silencing (*Dicer1, Ago1*), proteasome components (*Psmc5, Psmb1, Psme1*), as well as alterations in cellular respiration and glycolysis (*Idh2, Idh3g, Idh3b, Mdh1*). Together, these gene programs may reflect alternative inflammatory environments found deep within the tumor and/or stress-related pathways associated with hypoxia and cell death. We also found that Monocytes/Macrophages within the tumor core upregulate *Xbp1*, which has been previously described in tumor-associated myeloid cells to promote tumor growth through cell non-autonomous inhibition of anti-tumor lymphocytes^27^. Together, application of SPACECAT to the KP lung tumor model recovered spatially-restricted cellular phenotypes that match prior knowledge of the diverse roles of tumor-associated macrophages, while also pointing to new transcriptional drivers of spatial context-specific immune responses.

## DISCUSSION

SPACECAT provides a broadly-applicable method for fluorescently tagging cells in arbitrary regions of interest, tracking of live cells over extended periods of time, and linking information about a cell’s spatial location or dynamics to myriad other measurements (e.g., single-cell -omic measurements, flow cytometry). Using *in vitro* cell cultures, we demonstrated cell tagging with spatially specific and temporally stable fluorescent labeling. In three-dimensional human organoids, we confirmed known biological structures of stem cell-enriched, mitotic crypt niches, and in spleen tissue sections, we targeted T cell rich zones and recovered the expected enrichment. To uncover new biological phenotypes, we further applied SPACECAT to cross-sections of an autochthonous lung tumor mouse model and uncovered spatially-enriched rare subsets of monocytes exhibiting transcriptional programs associated with differentiation and immune recruitment, an exciting implication for understanding the spatial structure underlying heterogeneity in cancer-immune crosstalk.

In this work, we have leveraged flow cytometry and scRNA-seq as final readouts after cell tagging using the SPACECAT protocol. Importantly, this approach yields live cells and allows for significant experimental flexibility. Cells can either remain *in situ* for dynamic tracking, fixation, and staining, or can be removed from that environment to be evaluated by biophysical assays, functional characterization, genomics methods, or potentially these assays in series. With the minimally perturbative and temporally stable fluorescent tagging signal, the SPACECAT protocol can be incorporated into existing sample workflows to increase information produced from an experiment, requiring only the addition of specific dyes described here and elsewhere^14^, and an active UV illumination source. By exposing a sample to different dyes, separated by washing and photoactivation steps, this protocol can yield tagged cells from multiple regions to be compared and contrasted. Importantly, the number of possible regions scales with the number of dyes, enabling many neighborhoods to be tagged from a limited set of encoding dyes.

By enhancing endpoint readouts—whether they be cell growth rate, cytokine secretion, genomics, etc.—with positional information (e.g., interfaces between diseased tissue and normal parenchyma, proximity to lymphatics or a mucosal surface) or by enabling cells of interest to be tracked through space and time without genetic modifications, SPACECAT has the potential to further our understanding of connections between microscale cellular characteristics, mesoscale intercellular architecture, and macroscale biological function.

## MATERIALS AND METHODS

### Calcein NVOC synthesis

#### 4’,5’-Dimethyl-3’6’-diacetoxyspiro[isobenzofuran-1(3H),9’-[9H]xanthen-3-one (compound 1)

This material was synthesized using an adaption of a procedure from Lippard^13^. Phthalic anhydride (16.7 g, 0.113 mol) and 2-methylresorcinol (24.9 g, 0.206 mol) were charged in an oven-dried 250 mL round-bottomed flask under nitrogen, and slowly heated with the use of a sand bath (thermocouple probe in bath = 170°C) until the solids were completely melted. Anhydrous zinc chloride (15.0 g, 0.110 mol, weighed in an argon-filled glove box) was added portion-wise against positive nitrogen pressure as the sand bath temperature was slowly raised to 230°C. After approximately 30 minutes (as indicated by the literature), no solidification had occurred, so the temperature was further raised to 250°C, at which point the reaction began releasing water vapor.

A vent needle was added, and the reaction was left at a thermocouple temperature of 250°C until it had solidified as a dark red solid. The flask was removed from the sand bath and was allowed to cool in air for one hour. The solid was then removed and pulverized with a mortar and pestle.

The resulting powder was suspended in 6N hydrochloric acid (150 mL) and heated to reflux for 30 minutes. The resulting solid product was collected by vacuum filtration, washed with water and air-dried with flowing compressed air and continual vacuum pressure. The solid obtained was carried on crude into the next step.

This crude solid was suspended in acetic anhydride (300 mL) in a round bottomed flask under nitrogen atmosphere and heated to a bath temperature of 100°C. Once at temperature, pyridine (20 mL) was slowly added down from the top of the attached reflux condenser, and the reaction was left to stir overnight. In the morning, TLC analysis indicated complete consumption of the starting material (as judged against a small sample of crude solid reserved from the first step). The reaction was allowed to cool until it was only slightly warm to the touch and was then poured onto approximately 300 g of ice to induce precipitation.

After the ice melted, the resulting solid was collected by filtration and washed with water. Recrystallization from acetone yielded 24.2 g (50% yield) of a creamy solid that produced colorless solutions upon dissolution.

^1^H NMR (401 MHz, Chloroform-*d*) δ 8.17-7.91 (m, 1H), 7.66 (dtd, *J* = 21.9, 7.4, 1.2 Hz, 2H), 7.22 (d, *J* = 7.5 Hz, 1H), 6.78 (d, *J* = 8.7 Hz, 2H), 6.68 (dd, *J* = 8.7, 0.7 Hz, 2H), 2.36 (s, 6H), 2.35 (s, 6H).

^13^C NMR (150.9 MHz, Chloroform-*d*) δ 168.8, 152.8, 150.4, 150.2, 135.1, 129.9, 126.4, 125.5, 124.2, 119.1, 117.7, 116.4, 82.6, 20.8, 9.5. HRMS (ESI) calculated for C_26_H_20_O_7_ [M+H]^+^: 445.1287. Found 445.1291. (**Supplementary Fig. 1**)

#### 4’,5’-Bis-(bromomethyl)-3’6’-diacetoxyspiro[isobenzofuran-1(3H),9’-[9H]xanthen-3-one (compound 2)

This material was synthesized by adapting a procedure from a Lippard patent^28^. The methyl fluorescein derivative prepared above (3.91 g, 8.80 mmol) and 1,3-dibromo-5,5-dimethylhydantoin (3.87 g, 13.5 mmol) were charged in a one-liter flask with a Schlenk adapter attached. Chlorobenzene (550 mL) was added, and the solution was degassed by freeze-pump-thaw. 1,1’-azobiscyclohexanecarbonitrile (180 mg, 0.74 mmol) and acetic acid (140 µL) were added against positive nitrogen pressure and the reaction was heated to 40°C for 72 hours, during which time the color changed from nearly clear, to a bromine-related orange, and finally back to yellow.

The chlorobenzene solution was transferred into a separatory funnel with the aid of additional toluene, and then rinsed with four portions (250 mL each) of water heated to between 50–75°C. The organic layers were dried over MgSO_4_ and concentrated to yield a crude solid, which was recrystallized from 9:1 toluene:EtOH. The resulting solid was collected by vacuum filtration and rinsed with a small amount of cold toluene. The filtrate solution was concentrated and recrystallized a second time from toluene/EtOH to yield a second crop of crystals. Both fractions were independently analyzed by ^1^H NMR before being combined (3.64g total, 68% yield) for storage and further analysis.

^1^H NMR (401 MHz, Chloroform-*d*) δ 8.12-7.99 (m, 1H), 7.70 (dtd, *J* = 23.8, 7.4, 1.2 Hz, 2H), 7.32-7.21 (overlapping with CDCl3 signal, 1H), 6.92 (d, *J* = 8.8 Hz, 2H), 6.83 (d, *J* = 8.8 Hz, 2H), 4.82 (s, 4H), 2.42 (s, 6H).

^13^C NMR (150.9 MHz, Chloroform-*d*) δ 168.7, 168.3, 152.1, 150.3, 149.3, 135.4, 130.3, 128.6, 126.1, 125.4, 124.2, 119.0, 118.8, 116.8, 81.3, 20.9, 20.3. HRMS (ESI) calculated for C_26_H_18_Br_2_O_7_ [M+H]^+^: 600.9498. Found 600.9492. (**Supplementary Fig. 2**)

#### Iminodiacetic acid bis(2-nitro-4,5-dimethoxy)benzyl ester trifluoroacetate salt (compound 3)

2-nitro-4,5-dimethoxybenzyl bromide from commercial suppliers was found to be relatively impure but was upgraded to sufficient purity by dissolving in dichloromethane and filtering through silica (eluted with additional pure dichloromethane) before use. This routinely converted dark brown commercial material into a pale yellow solid after concentration in vacuo.

*N*-Boc-iminodiacetic acid (184 mg, 0.79 mmol), purified 2-nitro-4,5-dimethoxybenzyl bromide (460 mg, 1.67 mmol) and cesium carbonate (542 mg, 1.66 mmol) were charged in an oven-dried 25 mL round-bottomed flask under nitrogen. Dry acetonitrile (4.6 mL) was added, and the solution was left to stir at room temperature for one hour. TLC analysis at that point indicated slow conversion, and so the reaction was heated to 50°C overnight (approximately 16 hours).

In the morning, TLC showed trace residual bromide (as it was added in slight excess) and one major new UV-active product. The reaction solution was transferred to a separatory funnel and diluted with saturated aqueous sodium carbonate, which was then extracted thrice with dichloromethane. The combined organic extracts were dried over MgSO_4_, filtered, and concentrated in vacuo.

To remove excess bromide, the crude product was re-dissolved in a small amount of dichloromethane and loaded onto a silica plug. Elution with pure dichloromethane was performed until all excess bromide had been removed, and then further elution with 10% ethyl acetate in dichloromethane delivered the major new UV-active compound as a single spot.

After concentration in vacuo, the resulting intermediate was immediately re-dissolved in dry dichloromethane (4 mL), placed under nitrogen, and cooled to 0°C. Trifluoroacetic acid (1 mL) was slowly added, and the reaction was left to stir under nitrogen for four hours. TLC analysis at that point indicated complete consumption of the presumed intermediate. The reaction was allowed to warm to room temperature over the course of fifteen minutes, at which point approximately 15 mL of diethyl ether were added with stirring. This induced immediate precipitation of fine particles. Stirring was stopped and the particles were allowed to flocculate for ten minutes before vacuum filtration. The creamy white solid that was obtained was rinsed with a further portion of diethyl ether (approximately 10 mL), before being air dried for ten minutes. The creamy white solid was then transferred into a tared 20 mL scintillation vial and further dried under high vacuum to deliver the desired trifluoroacetate salt (458 mg, 91% yield).

^1^H NMR (401 MHz, Methanol-*d*_4_) δ 7.78 (s, 1H), 7.19 (s, 1H), 5.63 (s, 2H), 4.15 (s, 2H), 3.97 (s, 3H), 3.93 (s, 3H).

^13^C NMR (105.9 MHz, methanol-*d*_*4*_) δ 166.5, 153.7, 149.1, 140.4, 124.7, 111.8, 108.2, 64.6, 55.6. Two carbon peaks are unobserved and are likely found underneath the solvent residual. HRMS (ESI) calculated for [M]^+^ C_22_H_26_N_3_O_12_: 524.1511. Found 524.1513. (**Supplementary Fig. 3**)

#### Calcein NVOC (compound 4)

This material was synthesized by adapting a procedure from the patent literature^29^. A portion of the trifluoroacetate salt generated above was scooped directly into a separatory funnel, to which was added pure water, and then an equal portion of saturated sodium carbonate solution. This aqueous layer was shaken and extracted three times with dichloromethane, then dried over MgSO_4_, filtered, and concentrated into a tared 50 mL round bottomed flask. After concentration in vacuo and weighing, it was determined that 90 mg of the free base of Iminodiacetic acid bis(2-nitro-4,5-dimethoxy)benzyl ester had been delivered (0.17 mmol).

4’,5’-Bis-(bromomethyl)-3’6’-diacetoxyspiro[isobenzofuran-1(3*H*),9’-[9*H*]xanthen-3-one (26 mg, 0.043 mmol), sodium iodide (13 mg, 0.087 mmol), and 1,8 bis(dimethylamino)naphthalene (18 mg, 0.084 mmol; Proton-sponge®) were added to the round bottomed flask, which was then purged with nitrogen and an oven-dried reflux condenser was attached. Dry acetonitrile was added, and the reaction was heated to 75°C for 16 hours.

During the course of reaction, a fine red-orange solid precipitated, and TLC analysis indicated complete consumption of the bis(bromomethyl) starting material, and one major new UV active spot of nearly the same R_f_ as the amine starting material. The reaction solution was loaded directly onto a preparative TLC plate (1000 µm thickness), with care taken not to transfer any of the reddish precipitate onto the plate. The plate was eluted with 3:1 ethyl acetate:hexanes including 1% added triethylamine, and the new major spot collected by removing the silica from the plate. The collected silica was washed with 1% methanol in dichloromethane and filtered to yield the desired product as a thin yellow film (with a small amount of residual solvent) inside the round bottomed flask it was collected in (17.6 mg, 28% yield).

^1^H NMR (600 MHz, Acetonitrile-*d*_*3*_) δ 8.03-8.00 (m, 1H), 7.75-7.68 (m, 2H), 7.60 (s, 4H), 7.27-7.20 (m, 1H), 6.95 (s, 4H), 6.83 (d, *J* = 1.6 Hz, 4H), 5.31 (dd, *J* = 8.8, 0.7 Hz, 8H), 4.28 (d, *J* = 1.3 Hz, 4H), 3.85 (s, 12H), 3.79 (s, 12H), 3.78 (s, 8H), 2.22 (s, 6H).

^13^C NMR (105.9 MHz, acetonitrile-*d*_*3*_) δ 171.5, 170.2, 169.7, 154.7, 153.8, 153.0, 151.3, 140.8, 136.7, 131.3, 128.7, 127.3, 126.9, 126.0, 124.8, 120.2, 119.8, 117.7, 111.7, 109.2, 82.7, 63.8, 57.1, 56.9, 54.6, 47.6, 21.1. One carbon signal is observed, and is assumed to be under the solvent residual. HRMS (ESI) calculated for [M+H]^+^: 1487.3851. Found 1487.3861. (**Supplementary Fig. 4**)

### Cell culture

Murine NIH/3T3 cells (ATCC, CRL-1658) and human HEK293T cells (ATCC, CRL-11268) were cultured in Dulbecco’s modified Eagle’s medium (DMEM) with glutamate and supplemented with 10% fetal bovine serum (FBS) at 37°C and 5% CO_2_. Cells were passaged every two to three days, as well as on the day of experiments after photoactivation for flow sorting and downstream library preparation. To passage cells, media was aspirated, and the cells were rinsed with Phosphate-buffered saline without calcium chloride and magnesium chloride (PBS0, Life Technologies) prior to the addition of Trypsin-LE (Life Technologies). The cells were incubated with Trypsin-LE at 37°C for 5 minutes, followed by the addition of complete media. Cells were pelleted by centrifugation for 5 minutes at 300g at 4°C.

### Organoid culture

All studies were performed under protocols approved by the Massachusetts Institute of Technology (MIT) Committee on the Use of Humans as Experimental Subjects and the Institutional Review Board (IRB) protocols of Massachusetts General Hospital/Partners Healthcare. Organoids were cultured as previously described^30,31^. Small intestinal crypts were isolated as previously described from de-identified, IRB-approved human bulk surgical resections. Briefly, the bulk resection was washed with ice-cold Dulbecco’s PBS0 to clear the luminal contents. Next 2-4 mm pieces were removed from the epithelial surface with scissors and washed repeatedly by gently pipetting the fragments using a 10-ml pipette until the supernatant was clear. Fragments were rocked on ice with crypt isolation buffer (10 mM EDTA, Life Technologies, in PBS0 supplemented with 500X dilution of Primocin, Invivogen, 10 mM HEPES, Life Technologies, and 2% FCS, Life Technologies) for 30 min. After isolation buffer was removed, fragments were washed with cold PBS0 by pipetting up and down to release the crypts. Crypt-containing fractions were combined, passed through a 100-μm cell strainer (BD Bioscience), and centrifuged at 50g for 5 min. The pellet was resuspended in basal culture medium (2 mM GlutaMAX, Thermo Fisher Scientifc, 500X Primocin, and 10 mM HEPES in Advanced DMEM/F12, Life Technologies) and centrifuged at 50g for 5 min once more to remove single cells. Crypts were then cultured in a Matrigel culture system (described below) in small intestinal crypt medium (50X B27 supplement, Life Technologies, 1 mM N-acetyl-L-cysteine, Sigma-Aldrich, in basal culture medium) supplemented with differentiation factors at 37°C with 5% CO_2_. To plate, crypts were resuspended in basal culture medium at a 1:1 ratio with Corning™ Matrigel™ Membrane Matrix – GFR (Fisher Scientific) and plated at the center of each well of 24-well plates. Following Matrigel polymerization, 500 μl crypt culture medium containing growth factors EGF (50 ng/ml, Life Technologies), Noggin (100 ng/ml, PeproTech), FGF-2 (50 ng/ml, PeproTech), IGF-1 (100 ng/ml, PeproTech) and R-spondin 3 (500 ng/ml, PeproTech) and conditioned media supplement Afamin-Wnt3a (10X, MBL international) was added to each well. ROCK inhibitor Y-27632 (10 μM, R&D Systems) was added for the first two days of culture only. Cell culture medium was changed every other day. After 6 days of culture, organoids were passaged and differentiated. Briefly, culture gel and medium were homogenized via mechanical disruption and centrifuged at 300g for 3 min at 4°C. Supernatant was removed and the pellet resuspended in basal culture medium repeatedly until the cloudy Matrigel was almost gone. On the last repeat, the pellet was resuspended in basal culture medium, the number of organoids counted, and centrifuged at 300g for 3 min at 4°C. The cell pellet was resuspended in basal culture medium at a 1:1 ratio with Matrigel and plated at the center of each well of 24-well plates (∼250 organoids/well). Following Matrigel polymerization, 500 μl crypt culture medium minus growth factor EGF was added to each well. Cell culture medium was changed every other day. Following an additional 6 days of culture, organoids were used in SPACECAT imaging and staining.

### SPACECAT staining

On the day of experiments, aliquots of calcein NVOC (6 M in DMSO) were diluted 1:100 in PEG200 (Sigma-Aldrich), then in PBS to a final concentration of 1:5,000 for photoactivation of organoids and tissue sections, or 1:10,000 for photoactivation of *in vitro* cell lines. PA Janelia Fluor 549 (PA-JF549, Tocris) and PA Janelia Fluor 646 (PA-JF646, Tocris) were reconstituted and diluted according to manufacturer’s instructions.

Cells were stained for at least 30 minutes in an incubator at 37°C and 5% CO_2_. Prior to imaging and photoactivation of *in vitro* cultures, staining solution was exchanged for phenol-free, FBS-free DMEM, while for organoids and tissue sections, imaging and photoactivation occurred in the staining solution.

### SPACECAT photoactivation of *in vitro* cell lines

We implemented an Andor Mosaic 3 digital micromirror device on an IX83 Olympus Microscope to achieve the spatially patterned illumination required for user-directed region of interest tagging. The commercially available Mosaic 3 simultaneously illuminates of all pixels of all user-created ROIs within a single field of view, enabling rapid processing of many regions as compared with systems that raster laser light using galvanometer-based beam steering (e.g. Rapp Firefly or Andor Micropoint systems). Further, the Mosaic’s resolution is primarily limited by the choice of objective lens, enabling single-cell or even subcellular illumination if desired.

HEK293T cells were seeded in a glass bottom imaging plate (Eppendorf) two days prior to photoactivation. On the day of photoactivation, cells were stained with SPACECAT molecules (e.g., calcein NVOC for **Fig. 1c**; calcein NVOC, PA-JF549, PA-JF646 for **Fig. 1e**) as described in “SPACECAT Staining”. For characterization of SPACECAT’s spatial resolution and temporal stability (**Fig. 1e, 3a**), cells were also stained with CellTrace Calcein Red-Orange (ThermoFisher) and Hoechst (ThermoFisher) at 1:10^4^ and 1:10^5^ concentrations in Hank’s balance salt Solution (Life Technologies), respectively.

After staining, cells were transferred to a microscope stagetop incubator (Tokai Hit) for temperature, humidity, and CO_2_ control throughout photoactivation and imaging. Imaging was conducted on an Olympus IX83 inverted microscope through Olympus UPLSAPO objectives and Semrock bandpass filters, with illumination supplied by a SpectraX Lumencor LED source and images captured on a Hamamatsu ORCA FLASH 4.0LT CMOS camera. Cells were imaged to capture their baseline fluorescence, followed by photoactivation in arbitrary, user-specified regions using 0.5-10 second pulses of 405nm light controlled by a Mosaic 3 digital micromirror device (Andor). We determined that calcein NVOC was reliably uncaged following 10 seconds of photoactivation, while PA-JF549 and PA-JF646 dyes could be uncaged after 0.5 seconds. Subsequent imaging captured cells’ fluorescence levels after photoactivation.

### CellProfiler image analysis of photoactivation specificity and stability

CellProfiler^32^ was used to segment time course images of SPACECAT photoactivation and quantify the spatial precision and temporal stability of fluorescence signals. Hoechst images were used to identify cells’ nuclei under CellProfiler’s global threshold strategy and robust background thresholding method (lower outlier fraction = 0.05, upper outlier fraction = 0.05, threshold smoothing scale = 10, threshold correction factor = 1.0, suppress local maxima closer than 15 pixels). Cell boundaries were identified under CellProfiler’s Propagation method with Otsu adaptive thresholding of images into two classes (threshold smoothing scale = 10, threshold correction factor = 1, size of adaptive window = 120, regularization factor = 0.01). After segmenting images, each cells’ position within a given image and mean fluorescence over time were calculated, followed by plotting and visualization in MATLAB (Mathworks).

### SPACECAT photoactivation in co-culture environments

HEK293T and NIH/3T3 cells were seeded onto separate pieces of No. 1 coverglass two days prior to photoactivation. On the day of photoactivation, the pieces of coverglass were placed abutting each other on Nunc glass bottom dishes (ThermoFisher), and cells were incubated with calcein NVOC as described in “SPACECAT Staining”. Depending on the experimental condition, either HEK293T or NIH/3T3 cells within ∼3mm of the species interface region were photoactivated using 1 second pulses of 405nm light. As a negative control sample, neither species was photoactivated but was exposed to the calcein NVOC molecule. Cells were then lifted from the coverglass as described in “Cell Culture”, followed by flow sorting with a Sony SH800Z (gating on calcein NVOC fluorescence levels) into individual wells of a 96-well plate containing 5 μL of Buffer RLT (Qiagen) and 1% β-mercaptoethanol.

### Generation of scRNA-seq libraries using Smart-Seq2

scRNA-seq libraries were generated according to the SMART-Seq2 protocol^15,16^. Briefly, RNA from single-cell lysates was purified using AMPure RNA Clean Spri beads (Beckman Coulter) at a 2.2x volume ratio, and mixed with oligo-dT primer, dNTPs (NEB), and RNase inhibitor (Fisher Scientific) at 72°C for 3 minutes on a thermal cycler to anneal the 3’ primer to polyadenylated mRNA. Reverse transcription was carried out in a master mix of Maxima RNaseH-minus RT enzyme and buffer (Fisher Scientific), MgCl_2_ (Sigma), Betaine (Sigma), RNase inhibitor, and a 5’ template switch oligonucleotide using the following protocol: 42°C for 90 minutes, followed by 10 cycles of 50°C for 2 minutes, 42°C for 2 minutes, and followed by inactivation at 70°C for 15 minutes. Whole transcriptome amplification was achieved by addition of KAPA HiFi HotStart ReadyMix (Kapa Biosystems) and ISPCR primer to the reverse transcription product and amplification on a thermal cycler using the following protocol: 98°C for 3 minutes, followed by 21 cycles of 98°C for 15 seconds, 67°C for 20 seconds, 72°C for 6 minutes, followed by a final 5-minute extension at 72°C. Libraries were purified using AMPure XP SPRI beads at a volume ratio of 0.8x followed by 0.9x. Library size was assessed using a High-Sensitivity DNA chip (Agilent Bioanalyzer), confirming the expected size distribution of ∼1,000-2,000 bp. Tagmentation reactions were carried out with the Nextera XT DNA Sample Preparation Kit (Illumina) using 250 pg of cDNA per single cell as input, with modified manufacturer’s instructions as described. Libraries were purified twice with AMPure XP SPRI beads at a volume ratio of 0.9x, size distribution assessed using a High Sensitivity DNA chip (Agilent Bioanalyzer) and Qubit High-Sensitivity DNA kit (Invitrogen). Libraries were pooled and sequenced using NextSeq500/550 High Output v2 kits (75 cycles, Illumina) using 30-30 paired end sequencing with 8-mer dual indexing.

### Cell line co-culture data processing

Smart-Seq2 libraries cells were sequenced to a depth of 1.3 ± 0.06 million (mean ± standard error of the mean; SEM) reads per cell. Pooled libraries were demultiplexed using bcl2fastq (v2.17.1.14) with default settings, and aligned using STAR^33^ to the mouse UCSC genome reference (version mm10) and human UCSC genome reference (version hg19) simultaneously, and a gene expression matrix was generated using RSEM (v1.2.3) in paired-end mode. Cells with fewer than 100,000 aligned reads across either genome were eliminated from subsequent analysis. All analysis of gene expression was completed using the normalized RSEM output as transcripts per million (TPM).

### SPACECAT isolation of regions from human small intestinal organoids

On the day of photoactivation, organoids were harvested from Matrigel domes and excess Matrigel was removed from organoids through gentle pipetting with a wide-bore P1000 tip. Organoids were briefly spun down at 50g for 2min, with supernatant and excess matrigel carefully removed. Organoids gently resuspended with wide-bore P1000 tip and placed onto a glass bottom imaging plate (Eppendorf) and stained with calcein NVOC as described in “SPACECAT Staining”. Individual protruding crypts were selectively photoactivated with the Mosaic 3 (Andor) as described in “SPACECAT photoactivation of *in vitro* cell lines”, using 4 second pulses of 405nm light. Organoids were dissociated into a single-cell suspension through brief, but vigorous pipetting with a P1000 tip, followed by a 300g 5 min. pelleting and treatment with Trypsin-LE for 10 minutes, with vigorous pipetting at 5 minutes and 10 minutes. Single cells were again pelleted at 300g for 5 minutes, and then strained through a 30µm strainer to exclude clumps. Following dissociation, cells were flow sorted with a Sony SH800Z based on calcein NVOC fluorescence levels.

### SPACECAT isolation of regions from live tissue sections

All studies were performed under animal protocols approved by the Harvard Medical School and Massachusetts Institute of Technology (MIT) Committee on Animal Care and Institutional Animal Care and Use Committee (IACUC). Genetically engineered mice were generated by injecting low passage Kras^Lox-Stop-Lox-G12D/+^; p53^fl/fl^ (KP) embryonic stem cells from the C57BL/6 background into albino C57BL/6 blastocysts^23^. High degree chimeric mice were identified by coat color and were aged to 12 weeks prior to tumor initiation. Tumors were initiated by intratracheal instillation of 2 × 10^8^ plaque forming units of Adeno-SPC-Cre (University of Iowa) using previously established protocols. 16 weeks post tumor initiation, mice were euthanized by cervical dislocation and lungs were perfused by cardiac injection with PBS0 prior to removal from the thoracic cavity for fine dissection. Large tumors were identified upon close tissue examination and microdissected with fine surgical tools before placing into RPMI + 10% FBS on ice until further processing.

Tissue samples from lung, lung tumor, small intestine, and spleen were sliced into ∼1mm sections using an acrylic tissue slicer (Braintree Scientific, Inc.). Brain samples were embedded in 4% low-molecular weight agarose in PBS0 and sectioned live using a vibratome (velocity 1mm/s, amplitude 0.9mm, frequency 65Hz, 150µm sections) into cold PBS0. After sectioning, tissue was stained with calcein NVOC as described in “SPACECAT Staining” for at least 30 minutes prior to imaging and photoactivation. Brain sections were additionally stained with Hoechst (ThermoFisher) according to manufacturer’s instructions, while mouse spleen samples were incubated with PE anti-mouse TCR β chain antibody (BioLegend) according to manufacturer’s instructions. Tissue sections were then imaged and photoactivated as described in “SPACECAT Photoactivation of *In Vitro* Cell Lines,” using 12 second pulses of 405nm light.

After photoactivating the spleen and lung tumor sections, samples were dissociated by mechanical and enzymatic methods using the gentleMACS Tissue Dissociation Kit (Miltenyi Biotec). Resultant cell suspensions were stained with calcein violet (ThermoFisher, 1:1,000 dilution in PBS0) as an orthogonal indicator of viability. For spleen sections, flow cytometry using a Sony SH800Z quantified cells positive for TCR β chain antibody, while for lung tumor sections, cells were sorted by gating on positive for both calcein violet (viability following dissociation) and calcein NVOC (presence in the region of interest). All experiments leveraged calcein NVOC stained without photoactivation negative control for determining proper gating. Tumor samples were sorted into 500µL RPMI + 10% FBS and kept on ice for subsequent single-cell RNA-seq.

### Generation of scRNA-Seq libraries using Seq-Well

Seq-Well was performed as previously described with changes noted below^18–20^. Briefly, a functionalized PDMS nanowell array was loaded with uniquely-barcoded mRNA capture beads (ChemGenes) and suspended in complete media for at least 20 minutes. 15,000 cells or all of what was isolated by FACS was deposited onto the top of each chip and settled into the wells by gravity. The array was gently washed four times with PBS0 and finally with RPMI, and sealed using a plasma-functionalized polycarbonate membrane with a pore size of 0.01µm. Seq-Well arrays were sealed in a dry 37°C oven for 40 minutes before being submerged in a lysis buffer containing guanidium thiocyanate (Sigma), 1mM EDTA, 1% beta-mercaptoethanol and 0.05% sarkosyl (Sigma) for 20 minutes at room temperature. Arrays were washed and then incubated in hybridization buffer containing 2M NaCl (Fisher Scientific) with 8% (v/v) polyethylene glycol (PEG, Sigma) for 40 minutes at room temperature with gentle rocking. Afterwards, the mRNA capture beads with mRNA hybridized were collected from each Seq-Well array in wash buffer containing 2M NaCl, 3mM MgCl_2_, 20mM Tris-HCl, and 8% (v/v) PEG, and beads were resuspended in a master mix for reverse transcription containing Maxima H Minus Reverse Transcriptase and buffer, dNTPs, RNase inhibitor, a 5’ template switch oligonucleotide, and PEG for 30 minutes at room temperature, and overnight at 52°C with end-over-end rotation. Exonuclease digestion was carried out as described previously, then beads were washed with TE with 0.01% Tween-20 (TE-TW, Fisher Scientific) and TE with 0.5% SDS (Sigma), denatured while rotating for 5 minutes in 0.2 M NaOH, and resuspended in ExoI (NEB) for 1 hour at 37°C with end-over-end rotation^20^. Next, beads were washed with TE-TW, and second strand synthesis was carried out by resuspending beads in a master mix containing Klenow Fragment (NEB), dNTPs, PEG, and the dN-SMRT oligonucleotide to enable random priming off of the beads. PCR was carried out as described using 2X KAPA HiFi Hotstart Readymix and ISPCR primer, and placed on a thermal cycler using the following protocol: 95°C for 3 minutes, followed by 4 cycles of 98°C for 20 seconds, 65°C for 45 seconds, 72°C for 3 minutes, followed by 12 cycles of 98°C for 20 seconds, 67°C for 20 seconds, 72°C for 3 minutes, followed by a final 5-minute extension at 72°C. Post-whole transcriptome amplification proceeded as described above for SMART-Seq2 libraries, with the following exceptions: AMPure XP SPRI bead cleanup occurred first at a 0.6 x volume ratio, followed by 0.8x. Library size was analyzed using an Agilent Tapestation hsD5000 kit, confirming the expected peak at ∼1,000bp, and absence of smaller peaks corresponding to primer. Libraries were quantified using Qubit High-Sensitivity DNA kit and prepared for Illumina sequencing using Nextera XT DNA Sample Preparation kit using 900pg of cDNA library as input to tagmentation reactions. Amplified final libraries were purified twice with AMPure XP SPRI beads as before, with a volume ratio of 0.6x followed by 0.8x. Libraries from 3 Seq-Well arrays were pooled and sequenced together using a NextSeq 500/550 High Output v2 kit (75 cycles) using a paired end read structure with custom read 1 primer (Seq-Well CR1P): read 1: 20 bases, read 2: 50 bases, read 1 index: 8 bases.

### Seq-Well data processing for organoids and KP lung tumors

Reads were aligned and processed according to the Drop-Seq Computational Protocol v2.0 (https://github.com/broadinstitute/Drop-seq). Briefly, reads were first demultiplexed according to index read 1 using bcl2fastq (v2.17.1.14) with default settings. Read 1 was split into the first 12 bases corresponding to the cell barcode (CB), and the 13-20^th^ bases, which encode the unique molecular identifier (UMI). CBs, UMIs, and read 2 sequences with low base quality were discarded, as were any that contained non-random sequences (e.g. primer sequences, poly-A tails). Following CB and UMI tagging, read 2 was aligned to the mouse genome (version mm10) for all KP lung tumor experiments, and to the human genome (version hg19) for all organoid experiments using STAR v2.5.2b with default parameters including “--limitOutSJcollapsed 1000000 --twopassMode Basic”. STAR alignments were merged to recover cell and molecular barcodes, and any sequences within hamming edit distance 1 were merged, as these likely originated from the same original sequence. Additional methods to correct for bead synthesis errors in the CB or UMI are detailed in the Drop-Seq Computational Protocol v2.0 (“DetectBeadSynthesisErrors” function). Digital gene expression matrices for each Seq-Well array were retained following quality filtering and UMI-correction, and further processed using the R language for Statistical Computing. Cells with fewer than 200 unique genes were removed from analysis.

### Seq-Well data analysis to identify cell types

Data from human organoid experiments and mouse KP lung tumors were processed similarly in R to visualize and identify cell types. Data was normalized and scaled using the Seurat R package (https://github.com/satija.lab/seurat)^34^: transforming the data to log_e_(UMI+1) and applying a scale factor of 10,000. We confirmed equivalent depth and cell quality across each of our arrays and the absence of major batch effects introduced by sequencing work-up day or other technical factors, and thus did not regress any batch-related covariates out of our data, including individual cell quality or mitochondrial gene expression. To identify major axes of variation within our data, we first examined only highly-variable genes across all cells, yielding approximately 1,000-3,000 variable genes with average expression > 0.1 log-normalized UMI across all cells. An approximate principal component analysis was applied to the cells to generate 100 principal components (PCs). Using the JackStraw function within Seurat, we identified significant PCs to be used for subsequent clustering and further dimensionality reduction^35^. Critically, we completed all of the following analysis over a range of variable gene cutoffs and principal components to ensure that our cell identification results were robust to parameter choice, data not shown.

### Seq-Well data analysis of human organoid samples

After pre-processing and filtering we recovered 266 cells from the Whole Organoid sample, and 187 cells from the Calcein NVOC+ region (genes/cell: 1616 ± 46, UMI/cell: 5016 ± 159, mean ± SEM) (**Supplementary Table 1**). For 2D visualization and cell type clustering of the human organoid data, we used the Barnes-Hut implementation of t-distributed stochastic neighbor embedding (t-SNE) with “perplexity” set to 20 and “max_iter” set to 10,000 (**Fig. 2c-e**). To identify clusters of transcriptionally-similar cells, we employed unsupervised clustering with the Louvain algorithm with the Jaccard correction^36^. Briefly, this method involves constructing a k-nearest neighbor graph over the Euclidean distance between cells in the PC reduced space, followed by a shared nearest neighbor (SNN)-based clustering and modularity optimization^37^. We implemented this using the FindClusters tool within the Seurat R package with default parameters and k.param set to 10 and resolution set to 0.3. We used the Seurat function FindAllMarkers to identify differentially-expressed genes upregulated within each cluster compared to all other cells in the dataset, and tested differential expression using the likelihood-ratio test for single-cell gene expression (by setting “test.use” to “bimod”)^38^. The significantly differentially-expressed genes (using a Bonferroni-corrected p-value cutoff of 0.05) for each cluster were analyzed to attribute likely identities of cells in each cluster. Analysis of cell cycle stage was carried out using the Seurat function “CellCycleScoring” with default parameters (**Fig. 2e, 2f**). Statistical significance of proportional differences by cell cycle stage or cell type was carried out using a Chi-square test.

### Seq-Well data analysis of mouse KP lung tumors

After pre-processing and filtering we recovered 2,424 cells from Tumor 1, with 1,634 of these cells coming from the Whole Tumor, 686 from the Healthy/Tumor Border, and 104 cells from the Tumor Core. From Tumor 2, we recovered 2,023 total cells, 1,489 of which from the Whole Tumor, and 534 from the Healthy/Tumor Border (**Supplementary Table 2**, genes/cell: 819 ± 10, UMI/cell: 2238 ± 36, mean ± SEM). For 2D visualization and cell type clustering of the human organoid data, we used a Uniform Manifold Approximation and Projection (UMAP) dimensionality reduction technique (https://github.com/lmcinnes/umap) with “min_dist” set to 0.5 and “n_neighbors” set to 30 (**Fig. 4b, 4d, Supplementary Fig. 9a**). To identify clusters of transcriptionally-similar cells, we employed unsupervised clustering as described above using the FindClusters tool within the Seurat R package with default parameters and k.param set to 10 and resolution set to 0.5. Here, we intentionally underclustered our data to avoid erroneously splitting cells with shared cell type functions, as the variable genes calculated for this dimensionally-reduced space likely did not fully reflect more nuanced cell type differences (e.g. variable behavior between lymphocyte subtypes). Each cluster was sub-clustered to identify more granular cell types, requiring each cell type to express >25 significantly upregulated genes by differential expression test (FindMarkers implemented in Seurat, setting “test.use” to “bimod”, Bonferroni-adjusted p-value cutoff < 0.001). After assessment of each subcluster, we identified 20 unique cell types across both tumors (**Fig. 4b, 4c, Supplementary Table 3**) Hierarchical clustering of cell types was conducted using complete linkage over the Spearman correlation between each cell type, computed over differentially-expressed genes between each cell type and all other cell types (**Fig. 4c**). Differences in abundance of cell types by photoactivation region (Whole Tumor vs Healthy/Tumor Border) was assessed by t-test (**Fig. 4e, Supplementary Fig. 9b**).

Differential expression tests between cells from Whole Tumor samples vs. Healthy/Tumor Border samples within the Monocyte/Macrophage cluster were conducted using the likelihood-ratio test for single-cell gene expression as described above (**Fig. 4g-4i, Supplementary Fig. 9c**). To order Monocytes/Macrophages by their spatial distribution (**Fig. 4j**), we scored each cell by taking the sum of the 117 genes upregulated in Tumor 1 Monocytes/Macrophages found at the Healthy/Tumor Border, and ranked single cells by this score. To identify gene programs associated with this ranking, we calculated the Pearson correlation between all genes and the Healthy/Tumor Border score. We used permutation testing to assess correlation values for significance, resulting in 887 genes negatively associated with the Healthy/Tumor Border phenotype, and 577 genes positively associated with the Healthy/Tumor Border phenotype. Heatmap was generated using the heatmap.2 function within the gplots R package. Significantly enriched gene ontologies (**Fig. 4k**) were assessed using the Database for Annotation, Visualization and Integrated Discovery (DAVID)^39^. All genes and statistics resulting from differential expression tests are reported in **Supplementary Table 3**.

### Data and code availability

The raw data and gene expression matrices for all scRNA-seq data will be deposited in the Gene Expression Omnibus (https://www.ncbi.nlm.nih.gov/geo/). Digital gene expression matrices annotated with cell types, photoactivation regions, and other metadata can be found in **Supplementary Table 1** (human intestinal organoids, corresponding to **Figure 2**) and **Supplementary Table 2** (mouse KP lung tumors, corresponding to **Figure 4**. Gene lists corresponding to differential expression tests in **Fig. 4c, 4g-4k** and **Supplementary Fig. 9c** can be found in **Supplementary Table 3**. All R and MATLAB code for analysis available upon request.

## ACKNOWLEDGEMENTS

We thank Marko Vukovic, Siyi Huang, Miyeko Mana, and Rachael Danielle Brust for organoid cell culture, sharing sample tissue for protocol optimization, and instrument support. We thank the members of the Shalek lab for comments and discussion on the project. A.K.S. was supported by the Searle Scholars Program, the Beckman Young Investigator Program, the Pew-Stewart Scholars Program for Cancer Research, a Sloan Fellowship in Chemistry, the NIH (1DP2GM119419 and 2RM1HG006193), and the Ragon Institute of MGH, MIT and Harvard. C.G.K.Z. was supported by T32GM007753 from the National Institute of General Medical Sciences. C.N.T. is funded by the National Science Foundation (NSF) Graduate Research Fellowship Program. A.M.J. was supported by the Damon Runyon Cancer Research Foundation. Synthetic development, including reagents and instrument time, was supported by Start-Up Funds from MIT and University of Calgary.

## AUTHOR CONTRIBUTIONS

A.S.G., C.G.K.Z., C.N.T., T.J., J.F.V.H. and A.K.S. conceived the study. A.S.G., C.G.K.Z., and C.N.T. performed and analyzed *in vitro* experiments, microscopy and data. B.M., and A.M.J., under the guidance of T.J., conducted organoid and *in vivo* experiments, respectively. R.P.K., A.W.N., and J.F.V.H. performed organic synthesis and accompanying analysis. A.S.G., C.G.K.Z., C.N.T., J.F.V.H. and A.K.S. interpreted the results and wrote the manuscript with input from all authors.

## SUPPLEMENTARY FIGURE LEGENDS

**Supplementary Figure 1. NMR of molecule (1)**

^1^H NMR (**a**) and ^13^C NMR (**b**) of 4’,5’-Dimethyl-3’6’-diacetoxyspiro[isobenzofuran-1(3*H*),9’-[9*H*]xanthen-3-one (**1**).

**Supplementary Figure 2. NMR of molecule (2)**

^1^H NMR (**a**) and ^13^C NMR (**b**) of 4’,5’-Bis-(bromomethyl)-3’6’-diacetoxyspiro[isobenzofuran-1(3*H*),9’-[9*H*]xanthen-3-one (**2**).

**Supplementary Figure 3. NMR of molecule (3)**

^1^H NMR (**a**) and ^13^C NMR (**b**) of Iminodiacetic acid bis(2-nitro-4,5-dimethoxy)benzyl ester trifluoroacetate salt (**3**).

**Supplementary Figure 4. NMR of molecule (4)**

^1^H NMR (**a**) and ^13^C NMR (**b**) of Calcein NVOC (**4**).

**Supplementary Figure 5. Overview of SPACECAT applications and chemistry.**

**a.** Schematic: application of SPACECAT to record spatial information as fluorescence signals in an arbitrary region of interest in complex primary tissue. **b.** Chemical structure of calcein NVOC and standard calcein AM and their respective intracellular transformations. **c.** Distributions of fluorescence changes at each timepoint, for cells outside of photoactivated regions (grey, n = 1,347 cells across 8 fields of view) and inside photoactivated regions (green, n = 68 cells across 8 fields of view).

**Supplementary Figure 6. Multiplexing photo-activatable probes**

**a.** Representative images of color multiplexing by photoactivating arbitrary regions in HEK293T cells using sequential addition of 3 photoactivatable probes (calcein NVOC, PA-JF549, PA-JF646). By leveraging different photoactivation thresholds (10 seconds for calcein NVOC, 0.5 seconds for PA-JF549 and PA-JF646), we achieve 5-color encoding. Scale bar = 100μm. **b.-i.** HEK293T cells were incubated with different combinations of photoactivatable probes (rows) and ∼2-20% of all cells in each sample were exposed to near-UV light, inducing photouncaging or photoconversion. Cells were dissociated, exposed to a master mix of all 3 dyes (to account for background fluorescence from individual dyes), and analyzed by flow cytometry. Gates determined by cells in control sample well stained with all dyes without UV-excitation. Left: calcein NVOC vs. PA-JF646, middle: PA-JF549 vs. PA-JF646, right: calcein NVOC vs. PA-JF549. Numbers reflect percent of cells within each quadrant. **b.** photoactivated without photo-convertible dyes, **c.** photoactivated while exposed to calcein NVOC only, **d.** PA-JF549 only, **e.** PA-JF646 only, **f.** calcein NVOC and PA-JF549, **g.** calcein NVOC and PA-JF646, **h.** PA-JF549 and PA-JF646, **i.** calcein NVOC, PA-JF549, PA-JF646.

**Supplementary Figure 7. Testing SPACECAT precision by specific phototagging of co-cultured HEK293T and NIH/3T3 cells. a.** Schematic of modified species-mixing experiment to test precision of spatially-restricted photoactivation. Human (HEK293T) and mouse (NIH/3T3) cell lines were seeded on separate coverglass slides and placed abutting in an imaging dish with calcein NVOC. **b.** FACS analysis and sort gates. Three such dishes (and corresponding FACS samples) were created: (left) HEK293T cells photoactivated, (center) NIH/3T3 cells photoactivated, and (right) no photoactivation. **c.** SMART-Seq2 library alignments to dual mm10-hg19 reference genome. Points represent single cells plotted by number of reads aligned to mm10 (x-axis) or hg19 (y-axis). Red: alignments support mouse cell, blue: alignments support human cell, grey: indeterminate species by transcriptome alignment.

**Supplementary Figure 8. Application of SPACECAT for spatial tagging in tissue sections across health and disease.**

**a.** Representative images of mouse spleen samples stained with calcein NVOC and photoactivated in arbitrary regions (white dashed regions). Scale bars = 100µm. **b.** Cell percent viability by calcein violet staining after photoactivation. N=2 lung tumor samples per condition, p-value > 0.05 by two-way ANOVA with post-hoc pairwise tests. **c.** Photoactivation of Healthy/Tumor Border of KP lung Tumor 2. Scale bars = 500µm.

**Supplementary Figure 9. Applying SPACECAT to a KP autochthonous lung tumors uncovers spatial heterogeneity of immune infiltrate.**

**a.** UMAP embedding with points corresponding to cells only in Tumor 2. Points colored by sort condition, shades of red represent triplicate Seq-Well arrays from the Whole Tumor. Black points represent single cells photoactivated and sorted from the Healthy/Tumor Border. **b.** Proportion of sort condition across cell types. Red bars: Whole Tumor; black bars: Healthy/Tumor Border. Student’s T test, p-value <0.05 *. **c.** Volcano plot of differentially expressed genes between Monocyte/Macrophage cells by tumor region in Tumor 2.

**Supplementary Table 1**. Digital gene expression matrix and metadata for cells derived from human intestinal organoids.

**Supplementary Table 2**. Digital gene expression matrix and metadata for cells derived from mouse KP lung tumors.

**Supplementary Table 3**. Differentially expressed genes by cell type and tumor location in KP autochthonous lung tumors.

