## Supplementary Fig. 1 for "Spatially-Resolved Live Cell Tagging and Isolation Using Protected Photoactivatable Cell Dyes"

### Supplementary Figure 1

**a**

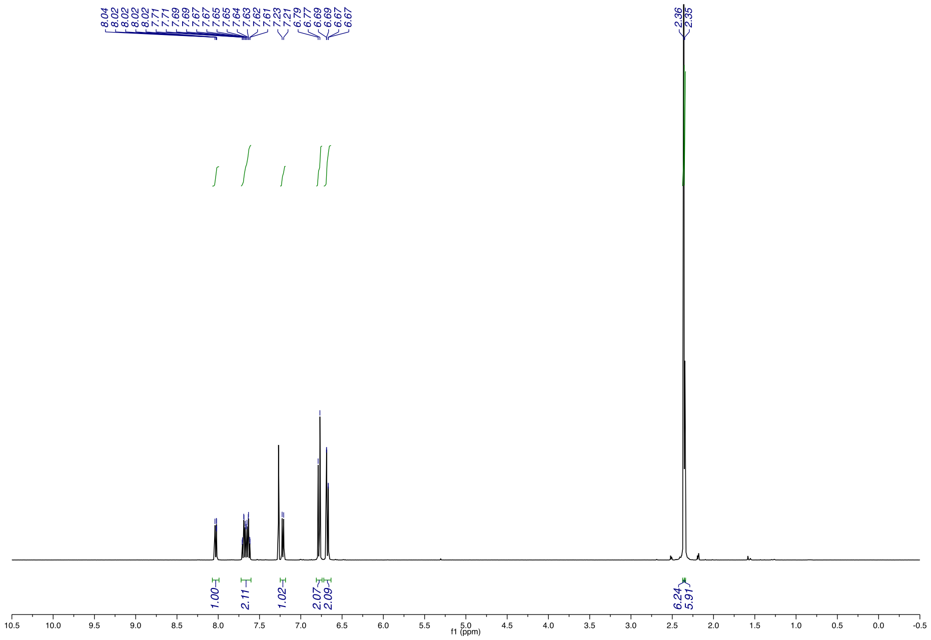


$^{1}$H NMR of 4',5'-Dimethyl-3'6'-diacetoxyspiro[isobenzofuran-1(3*H*),9'-[9*H*]xanthen-3-one (**1**)

**b**

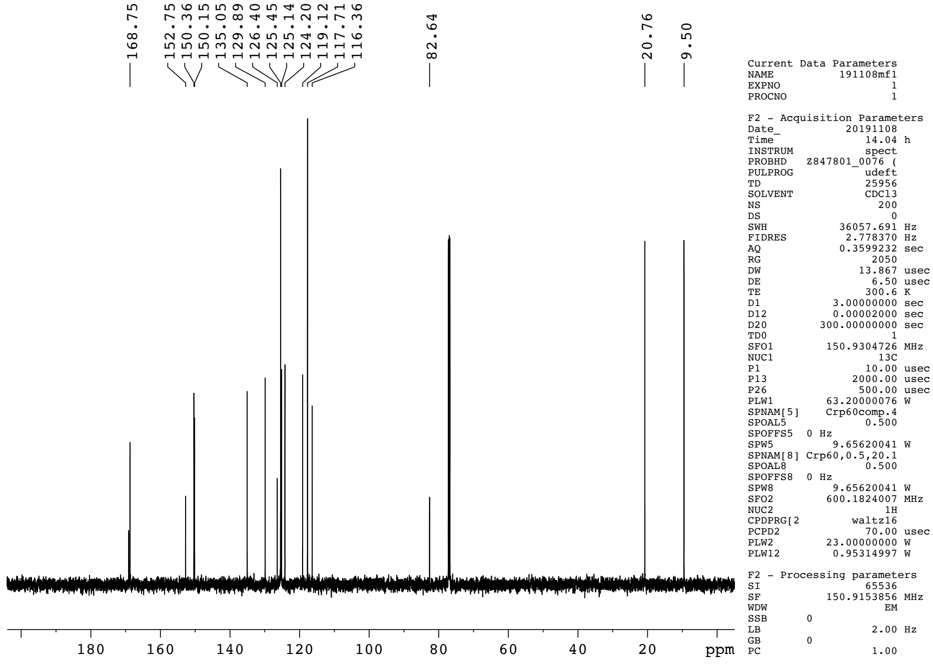


$^{13}$C NMR of 4',5'-Dimethyl-3'6'-diacetoxyspiro[isobenzofuran-1(3*H*),9'-[9*H*]xanthen-3-one (**1**)
