## Supplementary Fig. 2 for "Spatially-Resolved Live Cell Tagging and Isolation Using Protected Photoactivatable Cell Dyes"

### Supplementary Figure 2

## a

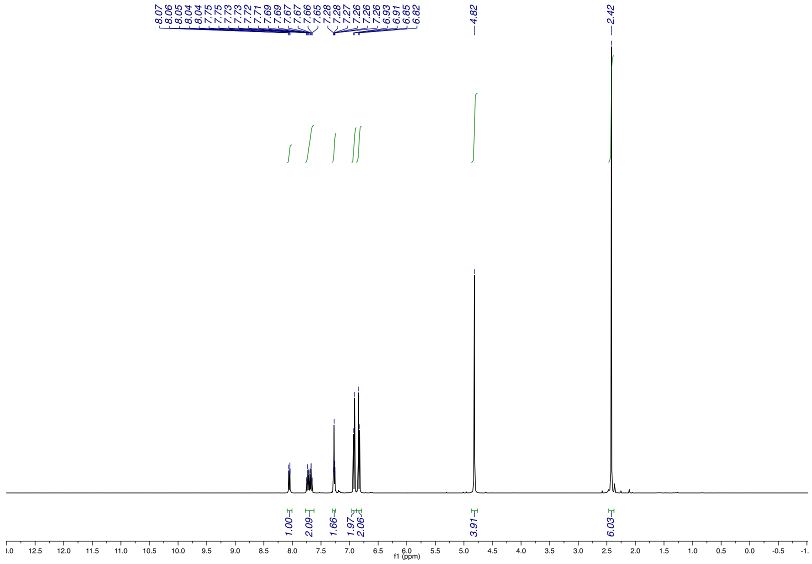


$^{1}$H NMR of
4’,5’-Bis-(bromomethyl)-3’6’-diacetoxyspiro[isobenzofuran-1(3*H*),9’-[9*H*]xanthen-3-one (**2**)

## b

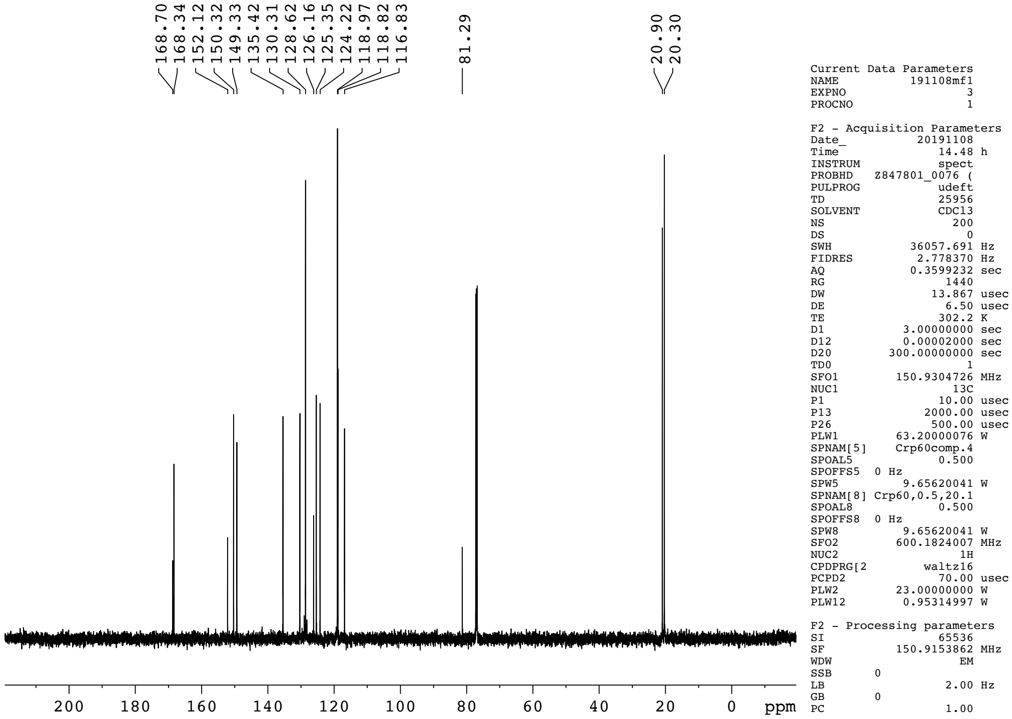


$^{13}$C NMR of
4’,5’-Bis-(bromomethyl)-3’6’-diacetoxyspiro[isobenzofuran-1(3*H*),9’-[9*H*]xanthen-3-one (**2**).
