## Supplementary Fig. 3 for "Spatially-Resolved Live Cell Tagging and Isolation Using Protected Photoactivatable Cell Dyes"

### Supplementary Figure 3

### a

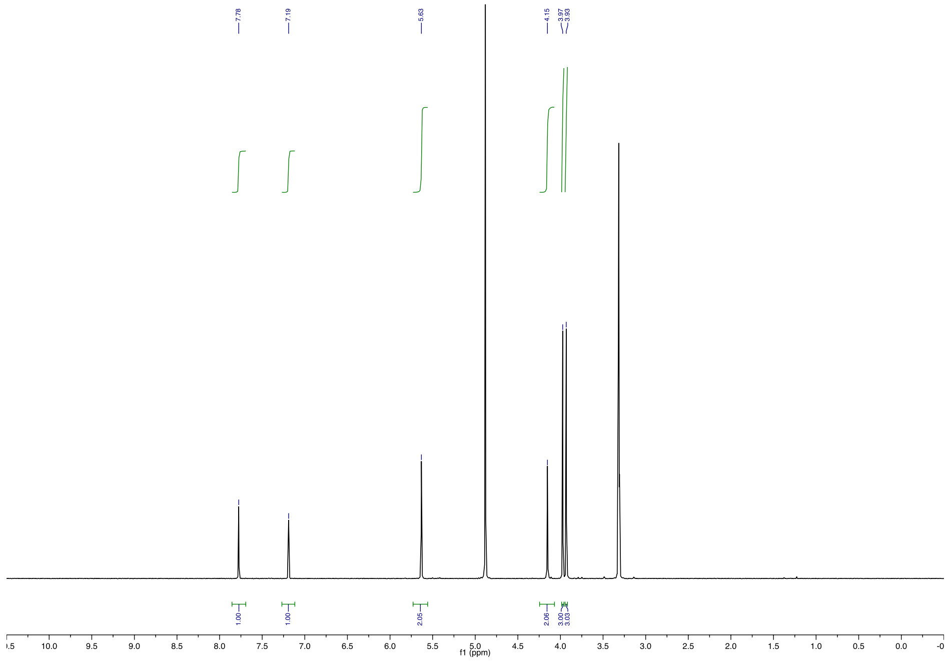


$^{1}H$ NMR of
*N*-Boc-iminodiacetic acid bis(2-nitro-4,5-dimethoxy)benzyl ester trifluoroacetate salt (**3**)

### b

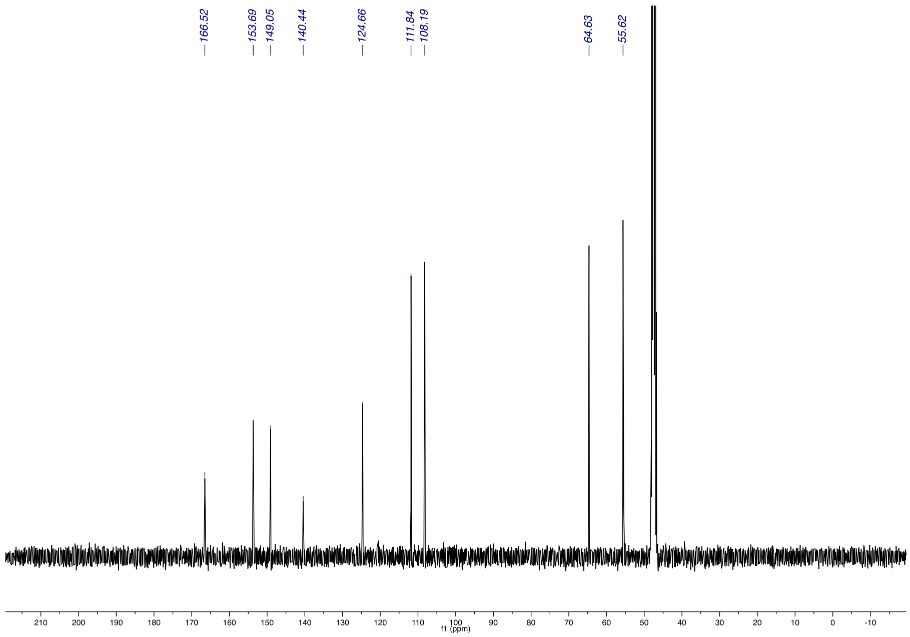


$^{13}C$ NMR of
*N*-Boc-iminodiacetic acid bis(2-nitro-4,5-dimethoxy)benzyl ester trifluoroacetate salt (**3**)
