## Supplementary figures and images for "Spatially-Resolved Live Cell Tagging and Isolation Using Protected Photoactivatable Cell Dyes"

### Supplementary Fig. 4

#### Supplementary Figure 4

**a**

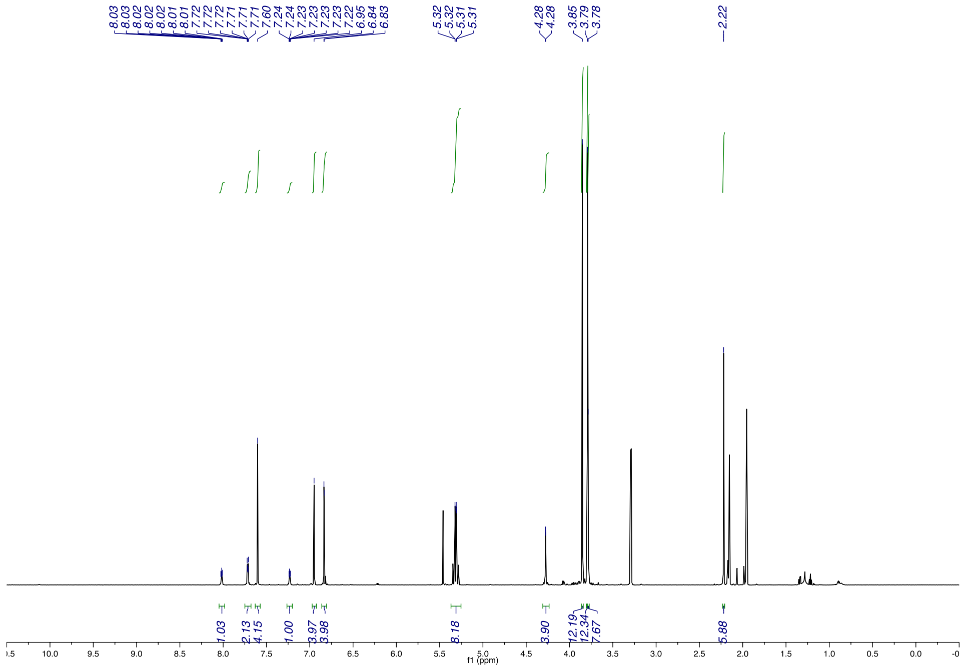

$^{1}H$ NMR of calcein NVOC (**4**)

**b**

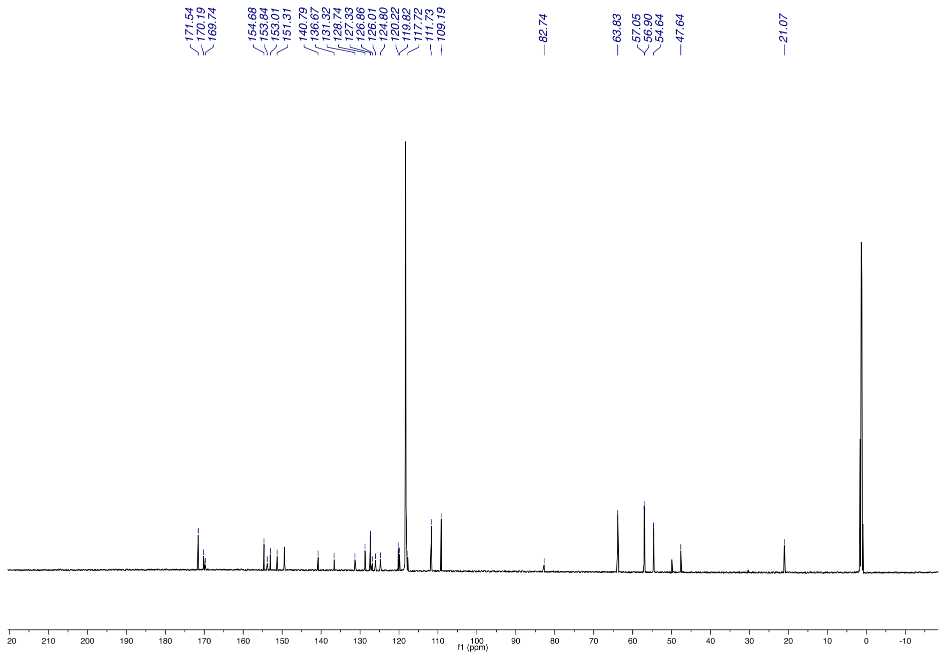

$^{13}C$ NMR of calcein NVOC (**4**)

### Supplementary Fig. 5

**Supplementary Figure 5**

a

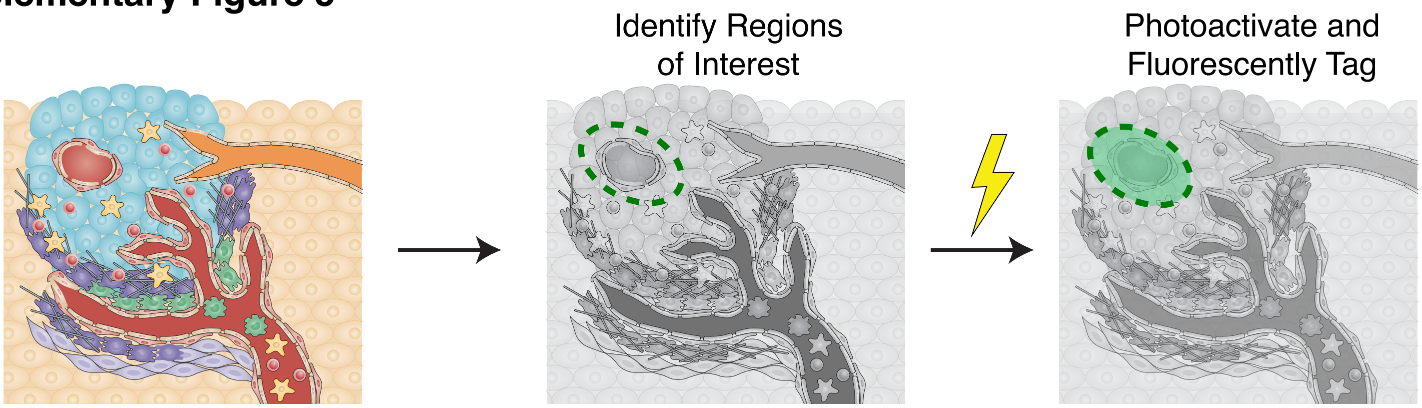


b

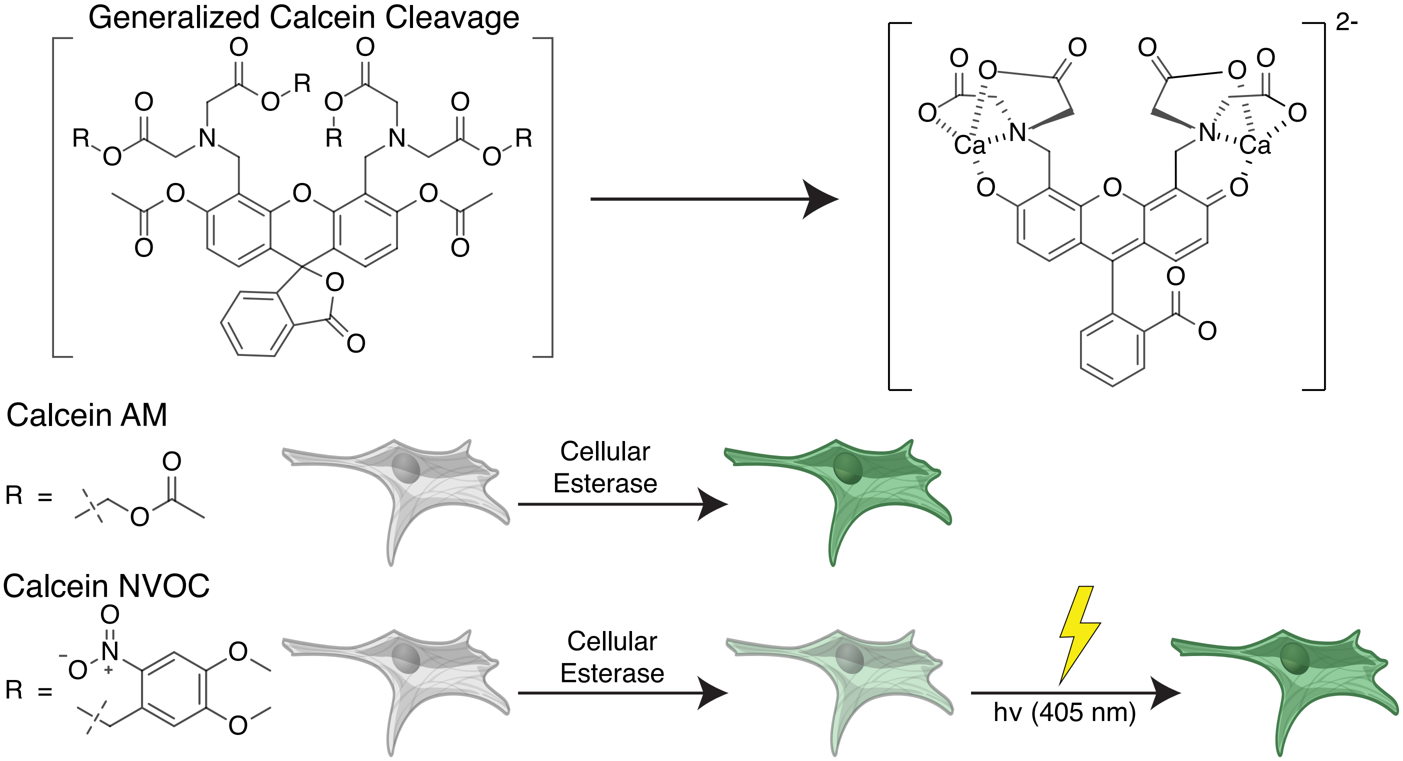


c

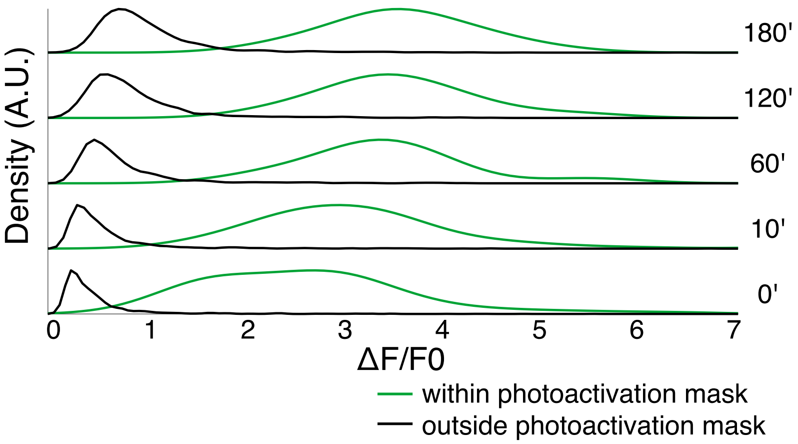

### Supplementary Fig. 6

#### Supplementary Figure 6

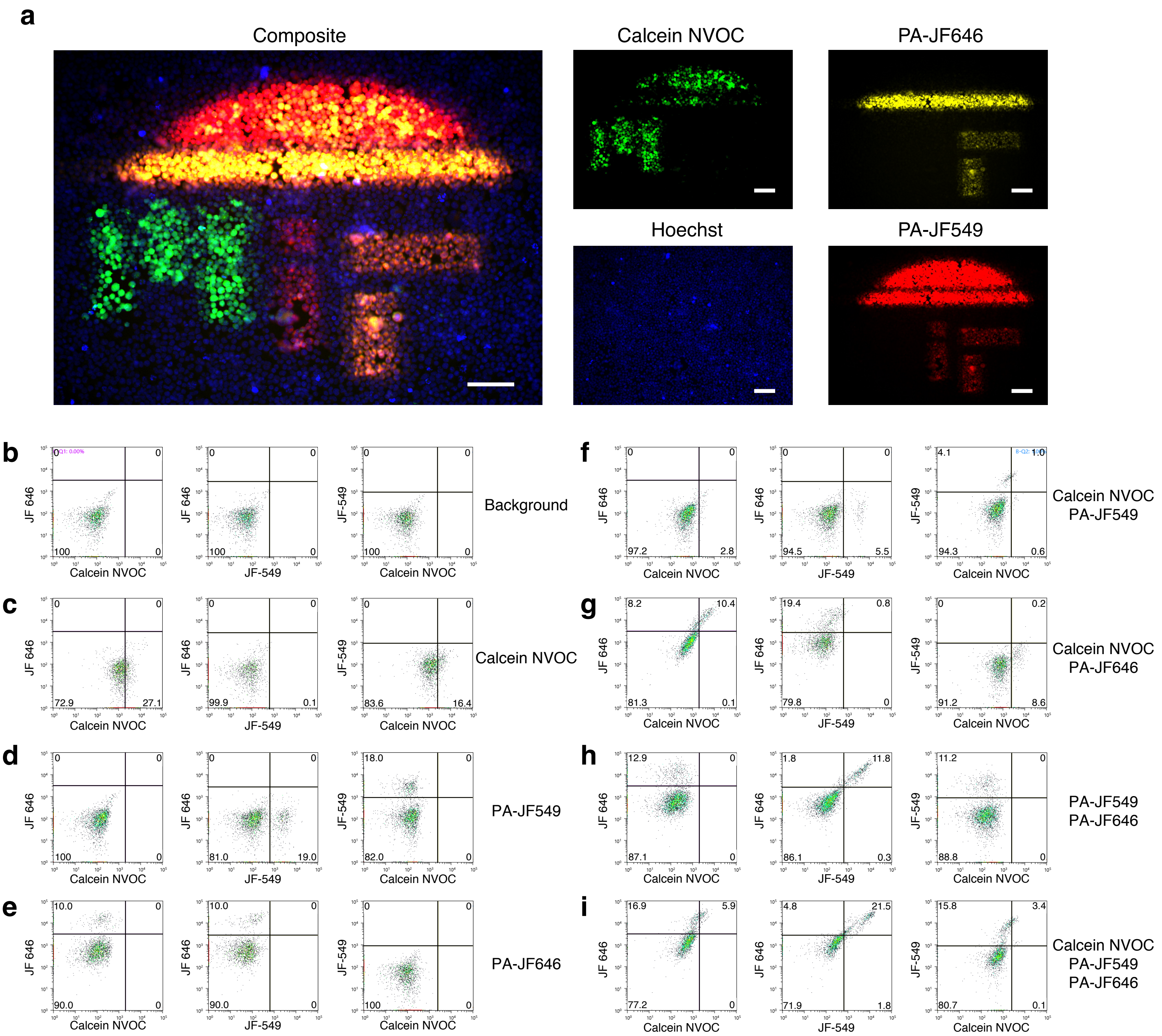

### Supplementary Fig. 7

#### Supplementary Figure 7

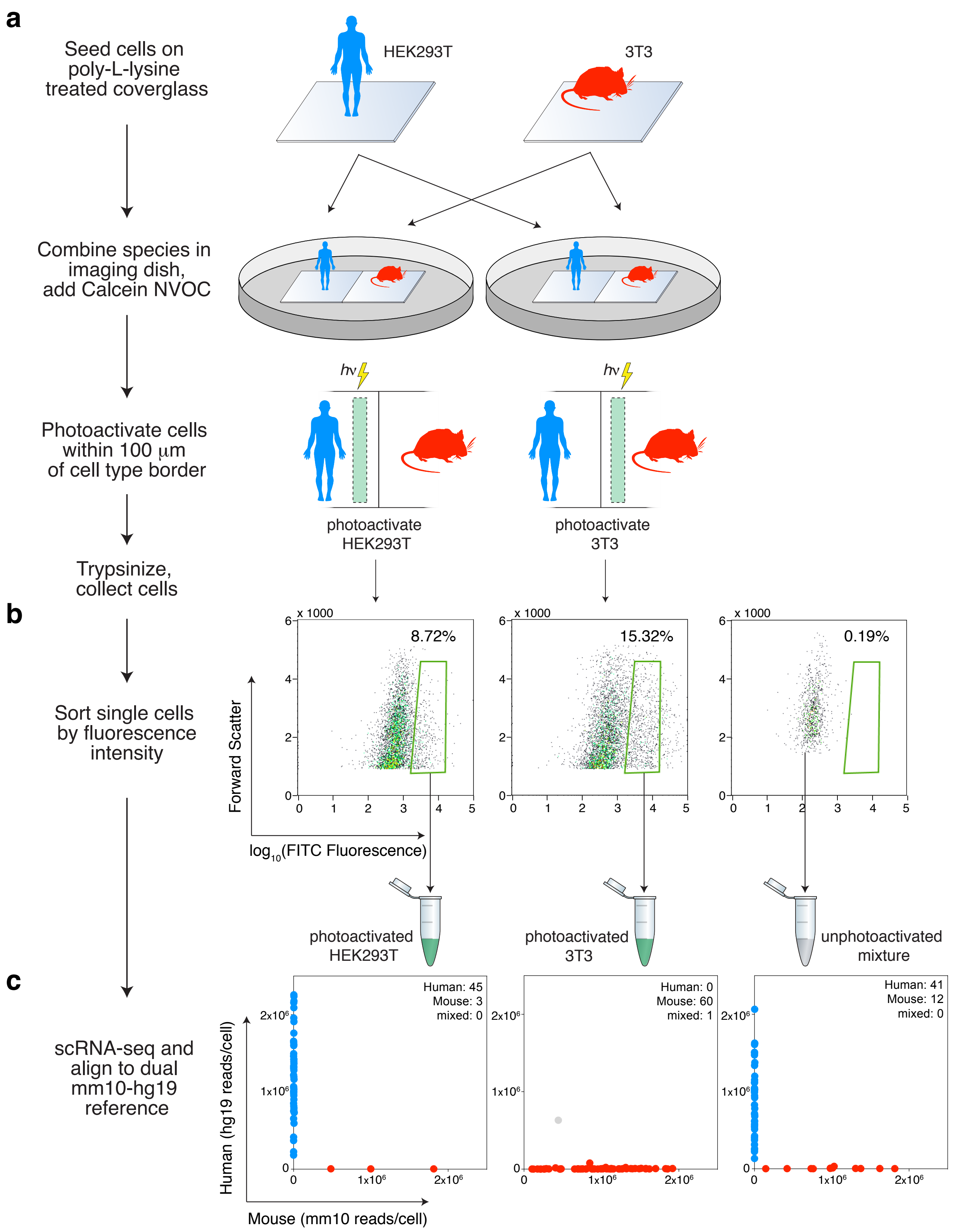

### Supplementary Fig. 8

#### Supplementary Figure 8

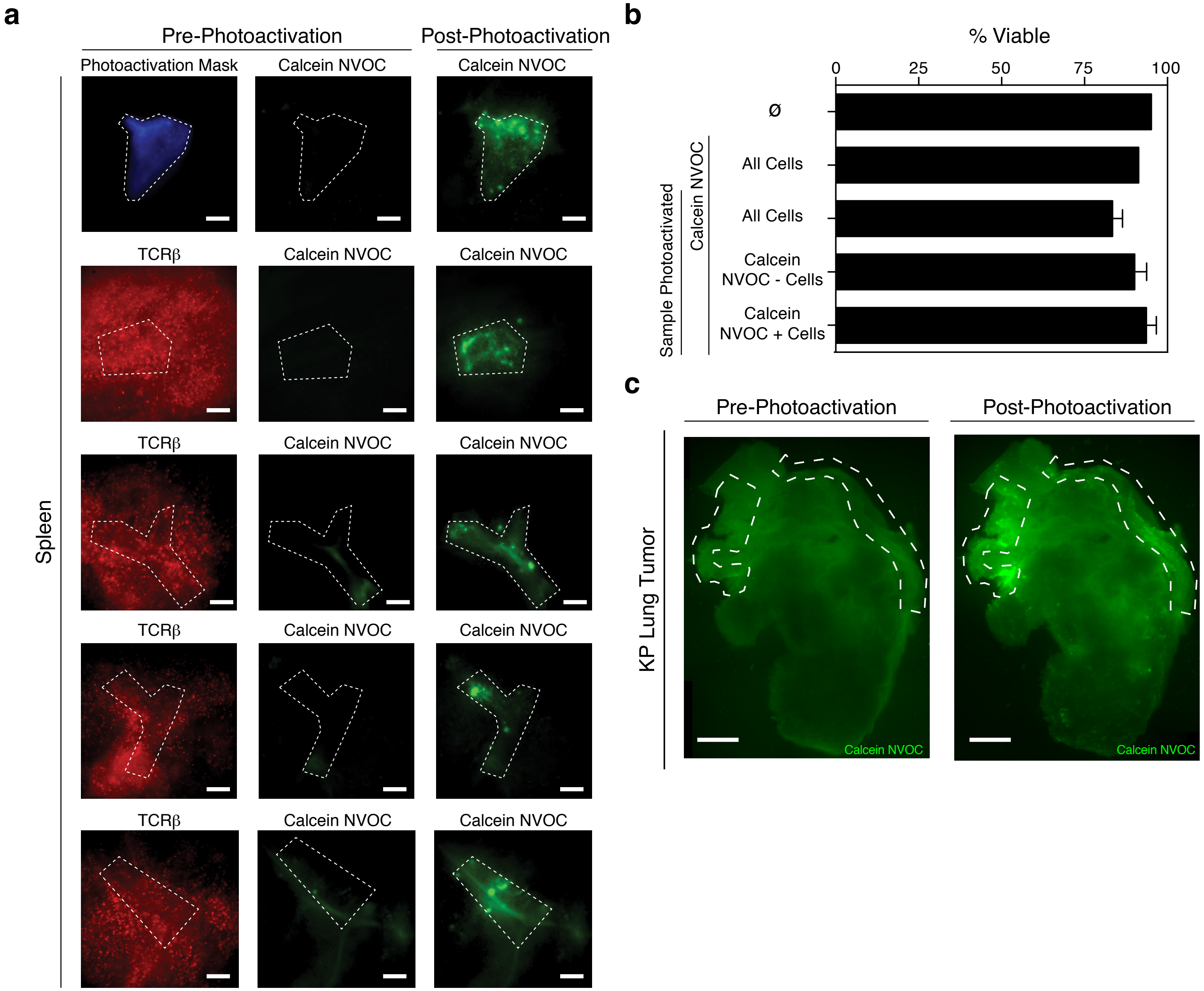

### Supplementary Fig. 9

#### Supplementary Figure 9

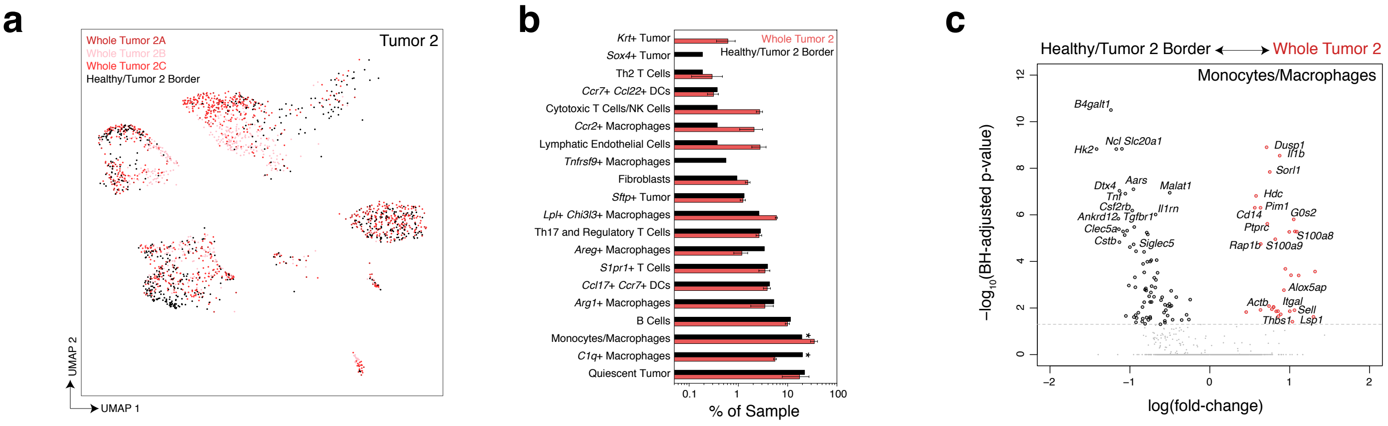
